# Variance (un)explained: Experimental conditions and temporal dependencies explain similarly small proportions of reaction time variability in linear models of perceptual and cognitive tasks

**DOI:** 10.1101/2022.12.22.521656

**Authors:** Marlou Nadine Perquin, Tobias Heed, Christoph Kayser

## Abstract

Any series of sensorimotor actions shows fluctuations in speed and accuracy from repetition to repetition, even when the sensory input and motor output requirements remain identical over time. Such fluctuations are particularly prominent in reaction time (RT) series from laboratory neurocognitive tasks. Despite their omnipresent nature, trial-to-trial fluctuations remain poorly understood. Here, we systematically analysed RT series from various neurocognitive tasks, quantifying how much of the total trial-to-trial RT variance can be explained with general linear models (GLMs) by three sources of variability that are frequently investigated in behavioural and neuroscientific research: 1) experimental conditions, employed to induce systematic patterns in variability, 2) short-term temporal dependencies as the autocorrelation between subsequent trials, and 3) long-term temporal trends over experimental blocks and sessions. Furthermore, we examined to what extent the explained variances by these sources are shared or unique. We analysed 1913 unique RT series from 30 different cognitive control and perception-based tasks. On average, the three sources together explained ∼8-17% of the total variance. The experimental conditions explained on average ∼2.5-3.5% but did not share explained variance with temporal dependencies. Thus, the largest part of the trial-to-trial fluctuations in RT remained unexplained by these three sources. Unexplained fluctuations may take on non-linear forms that are not picked up by GLMs. They may also be partially attributable to observable endogenous factors such as fluctuations in brain activity and bodily states. Still, some extent of randomness may be a feature of the neurobiological system rather than just nuisance.

**Public significance statement:** When we perform stereotyped repetitive sensorimotor actions, these nevertheless exhibit remarkable variability over time even under laboratory conditions. We show that most of this variability is not explained by linear models including two factors frequently studied in experimental psychology or cognitive neuroscience: the manipulations introduced by the experimenters and endogenous temporal dependencies that arise over multiple timescales. This implies that moment-to-moment fluctuations in typical behavioural data are largely unpredictable by linear models. Endogenous processes may also influence these fluctuations even in highly stable environments. It remains an empirical question to what extent exogenous and endogenous sources influence fluctuations non-linearly.

## Introduction

Any sensorimotor action we repeatedly execute, such as chopping vegetables, moving to a beat, or performing a defined move in sports, is characterised by variability in speed and accuracy from repetition to repetition. Such ***trial-to-trial variability*** is evident in typical laboratory tasks in which such sensorimotor actions are precisely recorded: although these tasks strictly control the environmental context with a limited set of sensory inputs and motor outputs, participants’ responses nevertheless fluctuate considerably from trial to trial. An example of this is shown in Figure 1 for the Flanker task (Figure 1A), a well-known paradigm for conflict processing. The series of RT measurements across trials for one participant (abbreviated as RT series hereafter) performing this task are highly variable (Figure 1B). Such fluctuations remain poorly understood: most of the time, we cannot explain why a participant responds fast or slow in a particular trial, and why they commit an error in one but not another. In other words, the predictability of individual actions over time appears highly limited. However, this predictability is important for empirical sciences in general (Yarkoni & Westfall, 2017), and also for daily life – for example, being able to predict why people sometimes respond fast (or slow) in critical tasks such as operating air control or driving a car (e.g., Baldwin et al., 2017). Here, we systematically examine trial-to-trial variability across a range of tasks that are commonly used in cognitive neuroscience – focusing on the linear trends as estimated by the general linear model (GLM). We concentrate on two well-documented constructs that both contribute to trial-to-trial variability, but which are typically studied separately: experimental conditions, which researchers may use to cause specific patterns in behavioural data, and task-unrelated fluctuations of behaviour over time. These temporal fluctuations may occur between subsequent trials (short-term) and across blocks of trials (longer-term). Here, we examine their contributions separately, allowing for an examination of three sources: experimental condition, short-term dependencies, and long-term trends.

**Figure 1.**
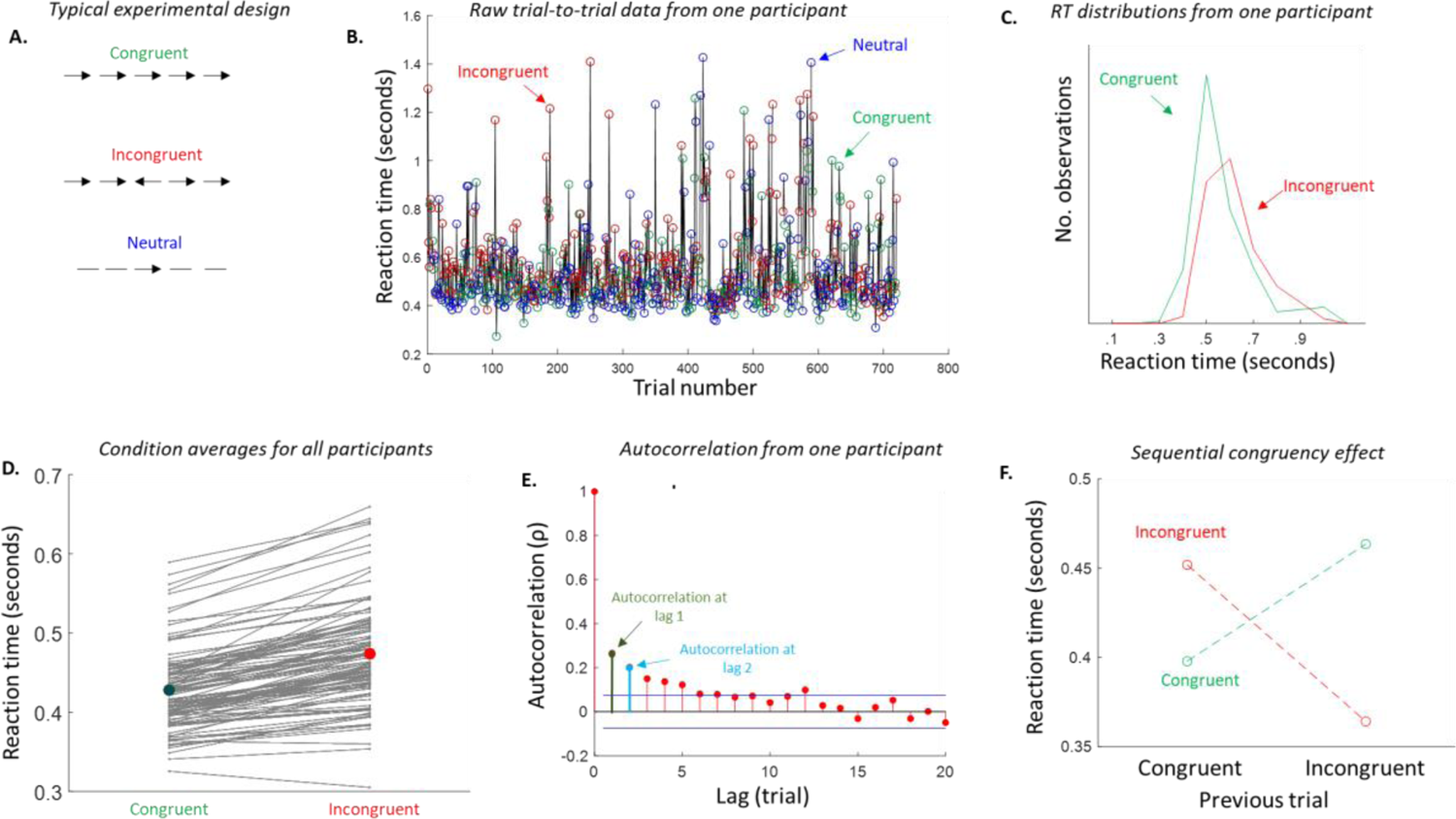
Example of a typical experimental study using the Flanker task. **A.** Participants discriminate the direction of the middle arrow while the flankers can be identical (congruent), contradictory (incongruent), or neutral. A researcher is examining whether participants are faster in the congruent and slower in the incongruent condition. **B.** Reaction time (RT) data from one participant on the Flanker task across all trials, based on data from Hedge et al. (2017). This reveals large fluctuations over trials across all experimental conditions (green, red, blue). **C.** RT distributions for the congruent and incongruent condition for one participant. The participant is faster in the congruent compared to the incongruent condition, but variability within each experimental condition is much larger than the average difference between experimental conditions. **D.** Means on the group level for congruent and incongruent conditions, with each line representing the RT of one participant. Nearly all participants are on average faster in the congruent than in the incongruent condition, in confirmation of the initial research question. **E.** Autocorrelation in RT series from one participant. Lag refers to the distance between trial numbers – e.g., lag one represents the autocorrelation between trial n and trial n-1. **F.** The effect of experimental conditions on RT in each trial is modulated by the experimental condition on the previous trial – a phenomenon known as the sequential congruency effect.

By implementing experimental conditions, researchers aim to induce systematic differences in behaviour to gain an understanding of specific neurocognitive mechanisms. However, systematic differences between conditions are often difficult to identify in individual data series. This is illustrated for the Flanker task example in Figure 1B: from the RT series, it is difficult to see whether the participant responds faster in congruent than incongruent trials. Fluctuations beyond those induced by the experimental conditions are usually considered ‘noise’ and are removed during data analysis by computing condition means or other forms of summary statistics. The experimental questions are then answered at the level of these summary statistics (e.g., “are participants *overall* slower on incongruent than on congruent trials?”). On this level, it is easier to identify patterns – for example, most participants are on average faster on the congruent than incongruent trials in the Flanker task (figure 1D), and the average across participants is visibly lower in the congruent condition. Visualising the distribution of reaction times (RT) for a single participant reveals high variance within conditions (Figure 1C), which is often much larger than variance between conditions. Such visualisations suggest that a large fraction of the within-subject variability in typical neurocognitive tasks is independent of the experimental conditions. In support of but not directly evidencing this claim, systematic group level analyses of cognitive conflict tasks show that the between-subject variance of the difference in mean RT between congruent and incongruent conditions is on average about seven times smaller than the within-subject RT variance (Rouder et al., 2019). However, most studies do not quantify or report what proportion of the within-subject variance is explained by the experimental effects (but see e.g., Williams et al., 2022). To address this question, the current study offers a systematic examination of the trial-to-trial variability that is explained by the experimental conditions across a range of tasks by GLMs.

Another line of research emphasises that the trial-by-trial variability of behaviour is not random but exhibits structured dependencies over time (e.g., Gilden, 2001; Van Orden et al., 2003; Perquin et al., 2023; Torre et al., 2019; Torre & Wagenmakers, 2009; Wagenmakers et al., 2004; Welford, 1968). For example, most RT series show positive autocorrelations, indicating that fast RTs are more likely followed by fast than slow RTs, and vice versa. Such positive autocorrelations are shown in Figure 1E for one participant in the Flanker task. Temporal dependencies have been found across many different tasks including cognitive control tasks, visual search, lexical decision, word naming, shape and colour discrimination, free choice RT, and racial implicit bias tasks (e.g., Annand & Holden, 2023; Correll, 2008; Gilden, 2001; Holden & Rajaraman, 2012; Marduski & LeBel, 2015; Van Order et al., 2003; Perquin et al., 2023; Simola et al., 2017). Importantly, they seem particularly present in tasks in which the same sensorimotor action is performed repeatedly without any changes in experimental condition, such as tapping to a fixed interval (e.g., Chen et al., 2002; Lemoine et al., 2006; Madison, 2001; Perquin et al., 2023). Because the external environment in such tasks remains unchanged, these dependencies must be driven endogenously – i.e., arising from fluctuations in the internal system. Such endogenous dependencies can emerge across multiple time scales and are important for the predictability of trial-to-trial fluctuations, as high dependency implies high predictability.

Certain forms of temporal dependencies can be caused by the order of experimental conditions and are hence induced exogenously (Duthoo et al., 2014; Egner, 2017; Fritsche et al., 2017; Gratton et al., 1992; Kayser & Kayser, 2018; Kiyonaga et al., 2017). An example is the ‘sequential congruency’ (alternatively, ‘Gratton’) effect, which denotes the phenomenon that participants respond faster if the previous trial was of the same condition as the current one (e.g., both congruent in the Flanker task; Figure 1F). Their interpretation is very different from endogenously-driven temporal effects: exogenously-driven effects reflect after-effects of external stimuli, while endogenously-driven effects may reflect temporal dynamics in spontaneous neural fluctuations (e.g., Palva et al., 2013; Smit et al, 2013). In the current study, we therefore included exogenous temporal effects in the form of experimental conditions, separately from short- and long-term endogenous temporal dependencies, allowing for a direct comparison of these sources of variability.

Here, we systematically examined the variance in the trial-to-trial variability of RTs explained by the experimental conditions and endogenous temporal trends across a sample of 1913 unique manual button press RT series. We focused on RT because it provides a continuous and rich measure of behavioural variability. The data were taken from publicly available resources and our own previous work and comprised tasks commonly used in the field of cognitive control (e.g., Flanker and Stroop tasks; termed ‘congruency-based’ in the following) and perception (e.g., detection of stimuli in noisy contexts and orientation discrimination; termed ‘perception-based’). The datasets are typical of experimental paradigms used in psychology and cognitive neuroscience: that is, the tasks are designed to study sensory and cognitive phenomena and are not specifically tailored to increase or reduce RT variability. We quantified how much of the total RT variability is explained by: 1) the experimental conditions, 2) short-range temporal dependencies between triplets of subsequent trials, and 3) long-range temporal dependencies across entire experimental blocks.

Datasets were analysed with GLMs, which is the standard model form to study neurocognitive effects in contemporary psychology and neuroscience (Blanca et al., 2018). We chose this model form as it allows for: 1) a consistent analysis pipeline across a large number of datasets – so that the presented results are comparable across all included datasets, and 2) for systematic comparison across the three sources – as non-linear models may dilute differences between them. However, the GLM only estimates within-subject variance that can be linearly explained by the three modelled sources. All presented results should be interpreted with these analysis choices in mind. We elaborate further on potential implications in the Discussion.

Changes in behaviour over the course of an experiment may or may not be related to changes in response to different conditions. For example, if a participant becomes slower over time due to increased drowsiness, it is possible that this affects difficult conditions (e.g., incongruent) more than easier ones (e.g., congruent), which would be reflected in an increase in the RT difference between conditions over time. Likewise, if a participant becomes better at responding to difficult but not to easier conditions with more practice, the difference between conditions decreases during the experiment. Both cases would be reflected in overlapping explained variance between experimental conditions and temporal dependencies. Indeed, in vigilance studies, responses to rare but not to frequent targets get slower and less accurate over the course of an experiment (Karimi-Rouzbahani et al., 2021; Temple et al., 2000), but it remains unclear whether such trends also exist in tasks in which the conditions are balanced and (pseudo-)randomly presented. Aside from experimental conditions, one might also expect common explained variance between short- and long-range temporal dependencies: for example, both dependencies could be characterisations of the same phenomenon, such as short-term autocorrelations producing longer trends. To address these questions of overlapping variance, we systematically examined to what extent the explained variances of the three sources are shared or unique using variance partitioning.

## Methods

Methods were the same for the congruency-based and the perception-based tasks, and are described for both below. Note that the labels ‘congruency-based’ and ‘perception-based’ are used as practical way to refer to the main field and theoretical interest of the task, not to a fundamental difference of task type. This study was not preregistered.

### Selection of datasets

We collected datasets from previously published studies (Figure 2), both from publicly available sources (e.g., OSF, github) and from our own work. We applied the following criteria to select datasets:

1. The experimental conditions had to be implemented in a within-subject design as a necessary condition for studying within-subject variance.
2. Experimental conditions were presented (pseudo-) randomly across trials (rather than in a blockwise manner), allowing us to investigate the relative contributions of the sequential and naturally-occuring temporal effects in experimental condition. For example, a task that requires only synchronous responses to a stimulus in one block and only asynchronous responses in the next block would not be selected.
3. The task was was not explicitly intended to affect participants’ performance over time; thus, for instance, learning or fatigue-inducing studies would not meet this criterion. Such paradigms experimentally induce temporal dependencies and are therefore not of interest in the current study, as we focus on endogenous temporal dependencies.
4. The task consisted of either one or two independent “experimental manipulations of interest” to facilitate analysis, because each additional manipulation would result in an increasingly disproportional amount of predictors for a GLM. Capturing two experimental manipulations A and B in the model requires six predictors (*A, B, A*B,* each on the current and previous trial), while three experimental conditions would require fourteen (2x *A, B, C, A*B, A*C, B*C, A*B*C*), and in analogy four experimental manipulations would require 28 predictors.
5. The task consisted of minimally 20 trials for each experimental manipulation of interest, autocorrelation lag, and block. For example, congruency-based task 1 (Table 1) has one experimental manipulation of interest (congruency), two autocorrelations, and 7 blocks of interest – which would require at least (10 * 20) 200 trials. Having sufficient observations per predictor is necessary for any regression model, and particularly important for the current within-subject GLMs as RT is a measure with high variability. Though there are no agreed-upon rules for the number of observations per predictor, we relied on a guideline that is considered stringent (Ogundimu et al., 2016).
6. The task featured at least 15 participants to obtain a reasonable estimate of the explained variance at the task level. We put the threshold for number of participants lower than for number of trials, because the analyses are run on the within-subject level.
7. The dataset includeed the raw (trial-to-trial) data and, for each trial, the RT, accuracy, experimental condition(s), trial, and block number (or, if missing, can be inferred from the original paper), and
8. RT were measured using manual button presses, the most commonly used response measure.

**Figure 2.**
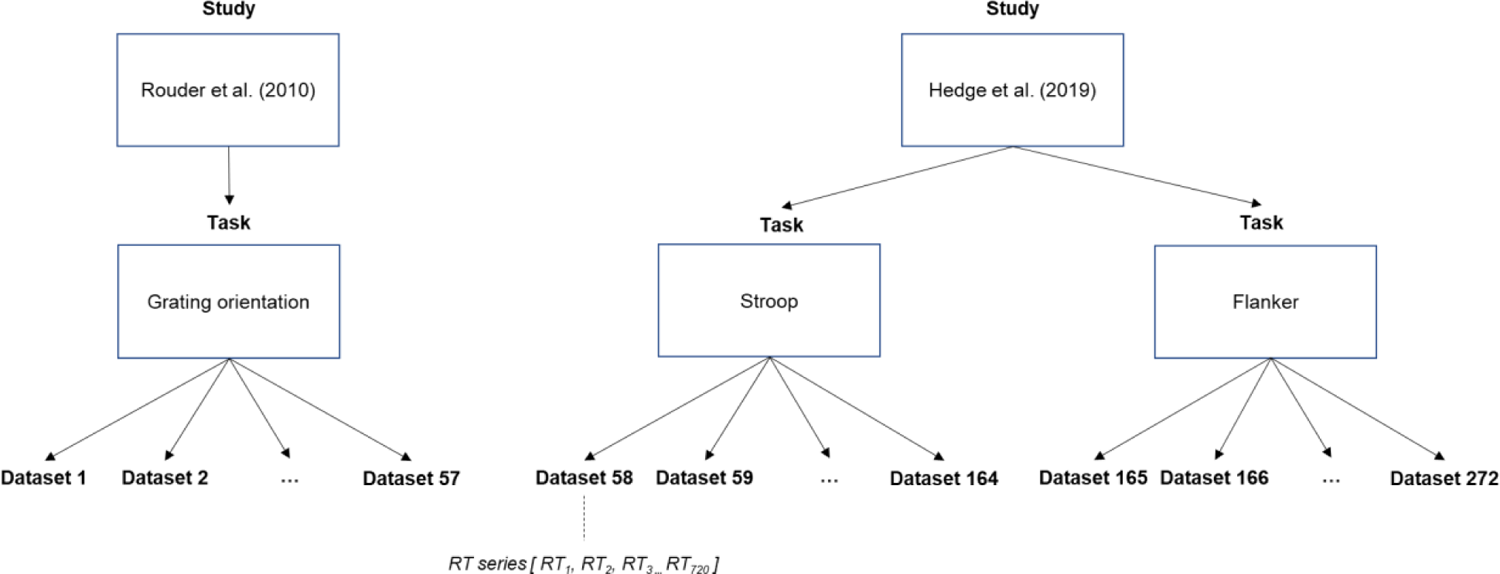
Overview of the nomenclature in the current research with the use of two examples. ‘Dataset’ refers to the trial-to-trial data from one participant on one task, including the RT, accuracy, and experimental condition for each trial. Datasets were sourced from previously conducted studies which contained one or more tasks. For example, we sourced 57 datasets from the grating orientation task from Rouder et al. (2005), and 214 datasets across two tasks from Hedge et al. (2019). All tasks with their original source and number of datasets are shown in Table 1 and 2.

**Table 1.** Overview of selected congruency-based task, showing the task name (task), total number of participants with complete data sets (# ppt), number of excluded participants in analysis based on % excluded trials (# excluded; see Methods above), the experimental manipulation(s) of interest with the number of conditions in brackets (# levels), the perceptual type of the stimulus (stimulus), the total number of blocks (blocks), the number of trials per block (trials/block) and per experiment (total trial #), the number of independent factors to predict RT (# predictors) for each participant, the original study in which the data was first published, and the original exclusion rules for RT outliers in the original study.

| **Dataset** | **#**<br>**datasets** | **#**<br>**excluded** | **manipulation**<br>**(# levels)** | **stimulus** | **blocks** | **trials/**<br>**block** | **Total**<br>**trial #** | **Original paper** | **Original**<br>**exclusion**<br>**rules** |
| --- | --- | --- | --- | --- | --- | --- | --- | --- | --- |
| 1. Stroop | 38 | 1 | congruency (3) | visual | 7 | 72 | 504 | Pratte et al.<br>(2010) | <200ms,<br>>2000ms |
| 2. Simon | 38 | 0 | congruency (3) | visual | 7 | 72 | 504 | Pratte et al.<br>(2010) | <200ms,<br>>2000ms |
| 3. Flanker | 107 | 4 | congruency (3) | visual | 5 | 144 | 720 | Hedge et al.<br>(2019) | <100ms,<br>>3SD+median |
| 4. Stroop | 107 | 1 | congruency (3) | visual | 5 | 144 | 720 | Hedge et al.<br>(2019) | <100ms,<br>>3SD+median |
| 5. Digit<br>distance | 54 | 3 | difficulty (3) | visual | 6 | 60 | 360 | Rouder et al.<br>(2005) | <250ms,<br>>2000ms |
| 6. Flanker<br>Arrow (young<br>participants) | 120 | 10 | congruency (3) | visual | 4 | 74 | 296 | Rey-Mermet et<br>al. (2018) | <250ms,<br>>2.5SD+mean |
| 7. Flanker<br>Arrow (older<br>participants) | 149 | 10 | congruency (3) | visual | 4 | 74 | 296 | Rey-Mermet et<br>al. (2018) | <250ms,<br>>2.5SD+mean |
| 8. Flanker<br>Letter<br>(young) | 120 | 13 | congruency (3) | visual | 4 | 74 | 296 | Rey-Mermet et<br>al. (2018) | <250ms,<br>>2.5SD+mean |
| 9. Flanker<br>Letter (old) | 148 | 18 | congruency (3) | visual | 4 | 74 | 296 | Rey-Mermet et<br>al. (2018) | <250ms,<br>>2.5SD+mean |
| 10. Stroop<br>Colour<br>(young) | 120 | 5 | congruency (3) | visual | 4 | 74 | 296 | Rey-Mermet et<br>al. (2018) | <250ms,<br>>2.5SD+mean |
| 11. Stroop<br>Colour (old) | 149 | 16 | congruency (3) | visual | 4 | 74 | 296 | Rey-Mermet et<br>al. (2018) | <250ms,<br>>2.5SD+mean |
| 12. Stroop<br>Number<br>(young) | 119 | 2 | congruency (3) | visual | 4 | 74 | 296 | Rey-Mermet et<br>al. (2018) | <250ms,<br>>2.5SD+mean |
| 13. Stroop<br>Number (old) | 147 | 17 | congruency (3) | visual | 4 | 74 | 296 | Rey-Mermet et<br>al. (2018) | <250ms,<br>>2.5SD+mean |
| 14. Priming<br>congruency | 31 | 4 | congruency (2) | visual | 8 | 80 | 640 | Desender et al.<br>(2016) | N/A |
| 15. Priming<br>congruency | 86 | 10 | congruency (2) | visual | 8 | 60 | 420 | Desender et al.<br>(2014) | none |
| 16. Priming<br>congruency | 30 | 4 | congruency (3) | visual | 6 | 144 | 846 | Teuchies et al.<br>(2019) | >1500ms |

We use the term ‘experimental manipulation of interest’ to refer to those manipulations that were of interest in the (statistical) comparison of the task at hand. For instance, in a Stroop task, experimenters use different words and colours for the stimuli, but the interest lies in the difference between congruent, neutral, and incongruent trials independently of the specific word-colour combination. As such, this task has one manipulation of interest (‘congruency’) that we use throughout the main analyses, as this is the driving source of experimentally induced variance. We will address the effect of other endogenous elements in the *Control analyses* (section *Additional sources of variance*).

### Selected datasets

#### Congruency-based datasets

Sixteen tasks were selected for analysis (see Table 1 for an overview, and the Supplementary Materials A for a full description). Of these, eleven were collected through a previous large-scale reanalysis of congruency-based tasks (Rouder, Kumat & Haaf, 2019), an additional three through the Confidence Database on OSF (Rahnev et al., 2019), and two were already in possession by the first author for another study (Perquin et al., 2023). We note that for these last two tasks (both from Hedge et al., 2018), the autocorrelation in RT has been previously reported alongside other temporal structure measures in Perquin et al. (2023) for a study of the reliability of temporal structures. All analysed datasets have been included in the current study.

#### Perception-based tasks

Fourteen tasks were selected for analysis (see Table 2 for an overview, and the Supplementary Materials A for a full description). Of these, six came from our own lab, one was found through the same large-scale reanalysis as above (Rouder et al., 2019), and an additional eight were found in the Confidence Database on OSF (Rahnev et al., 2020). One additional analysed task has not been included in the current results, as this concerned a simple detection task with a high number of anticipations, and all participants would have too little trials for the present purposes after data exclusion.

**Table 2.**
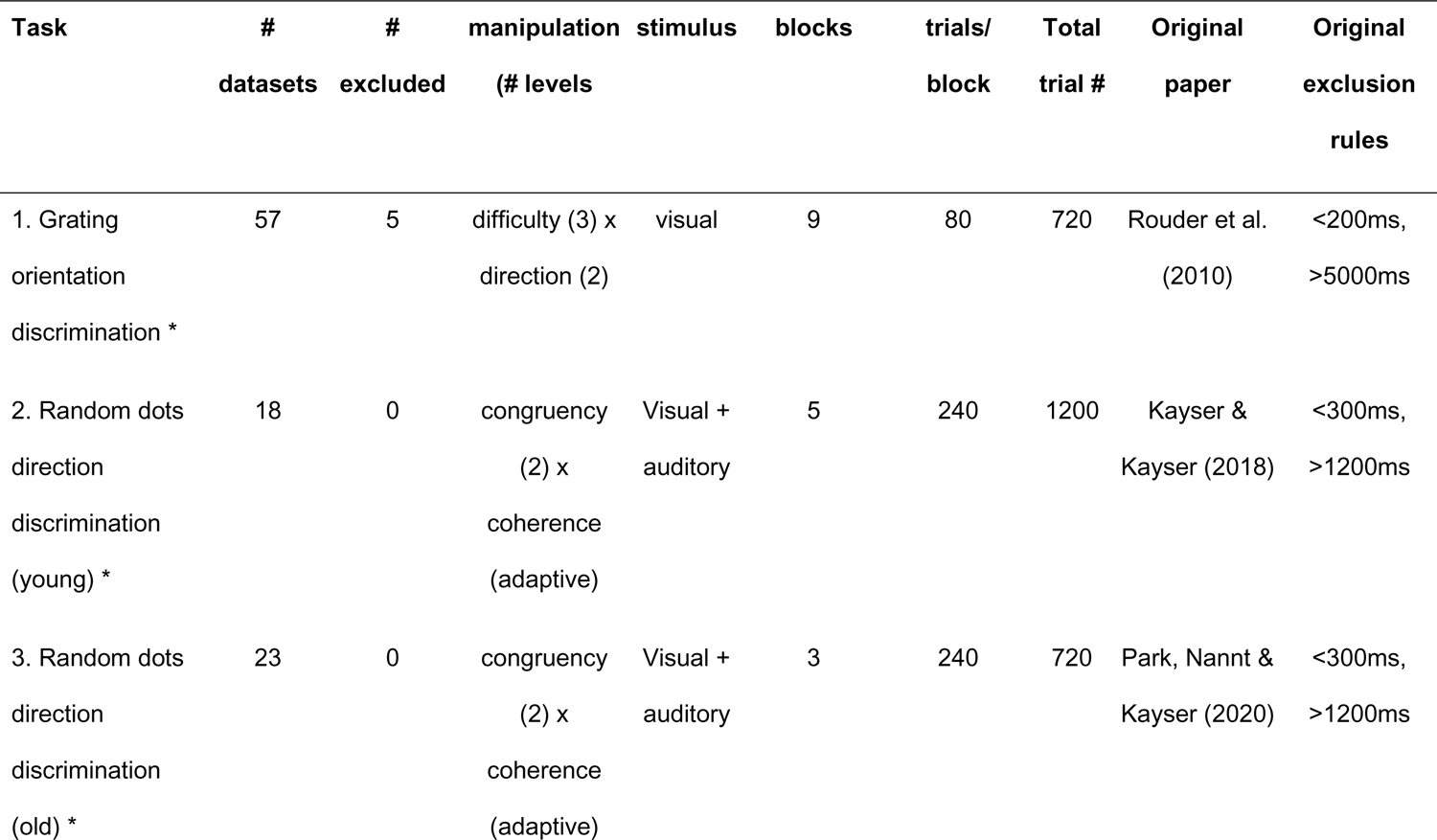
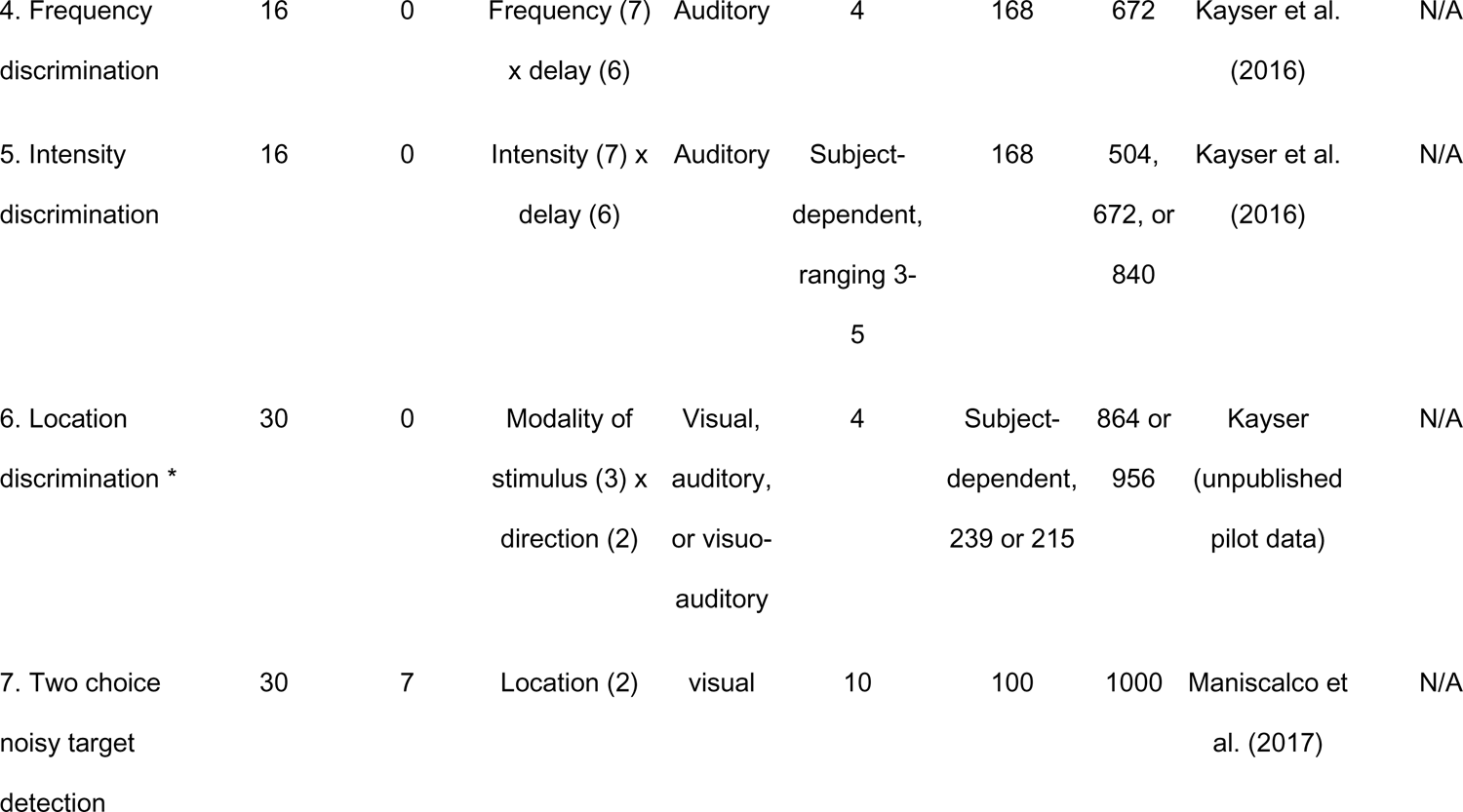
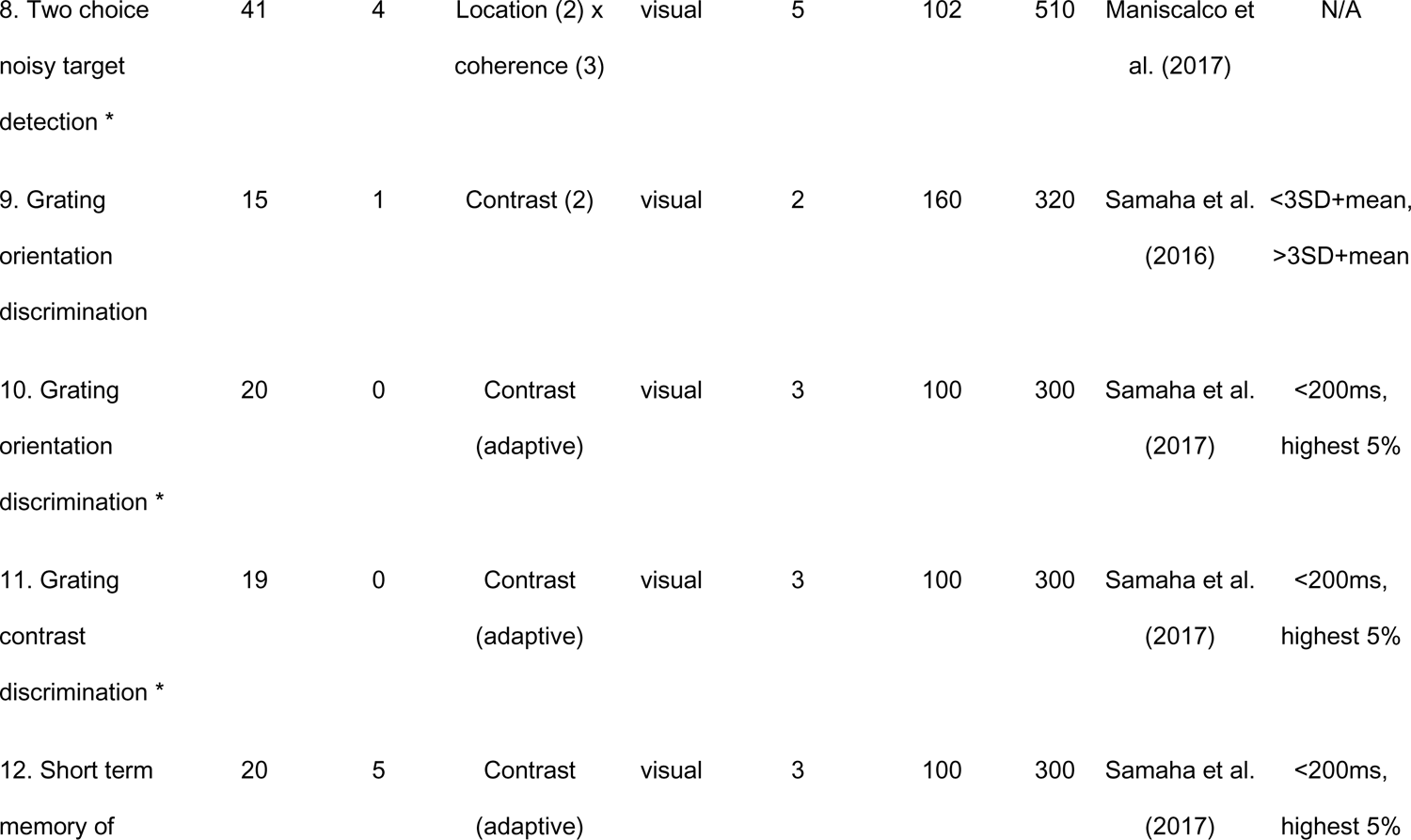
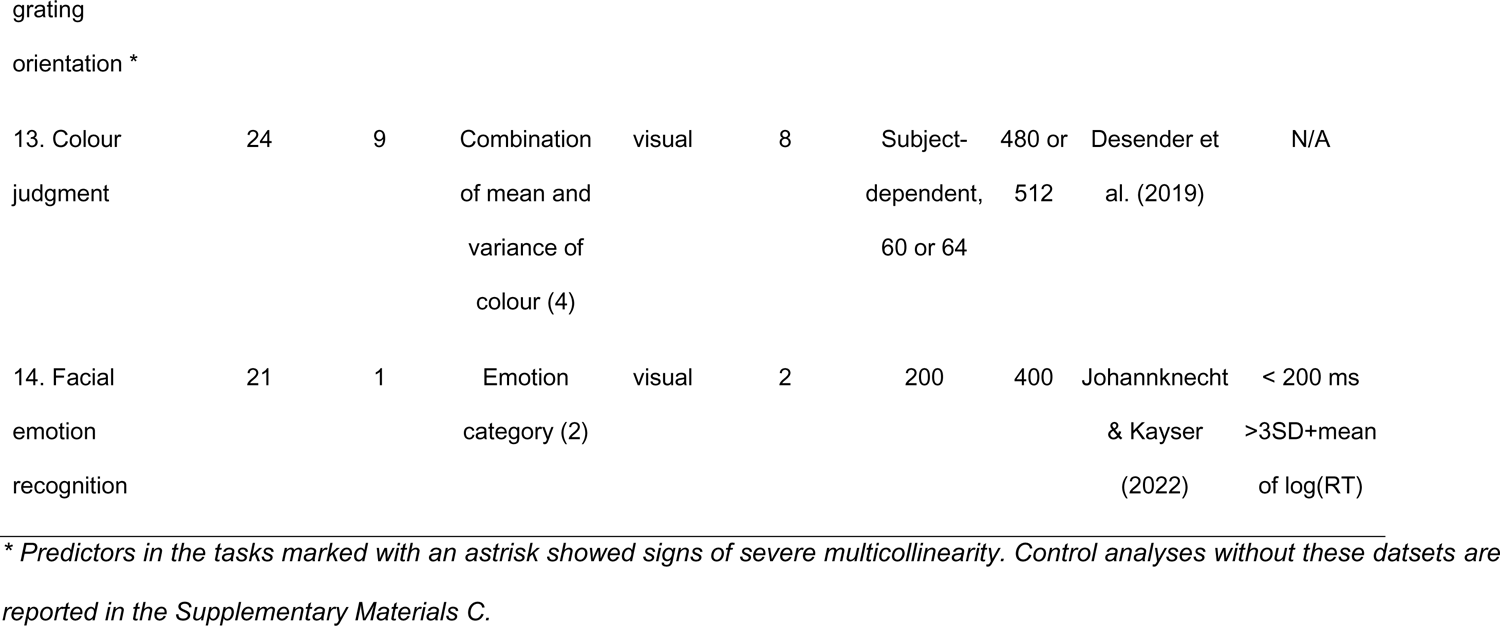
Overview of selected perception-based datasets. Same conventions as Table 1.

### Data preprocessing

Data cleaning was kept to a minimum to preserve within-subject fluctuations as faithfully as possible. Some cleaning is necessary: though the origins of ‘outliers’ remain largely unknown, they have a disproportionate weight on the fit of regression coefficients and were therefore excluded. First, excluded trials with RT > 10 s and RT < 150 ms were excluded under the assumption that these are not responses to the stimulus (but rather anticipations, experimenter errors, et cetera). Next, trials with RT > 3 standard deviations above the mean were excluded seperately for each participant and condition. The remaining RT series were inverse-transformed for all analyses to reduce their distributions’ skewness. The impact of these preprocessing choices on the overall results is discussed below (see section *Control analyses confirm that results are robust* in *Results*).

Trial with missing values were excluded from all analyses conducted in the current study. This seems to be the most common approach in studies on temporal dependencies, but it is important to note that the measurement of temporal dependencies is affected by trial exclusion (Karalunas et al., 2012; 2014; Kofler et al., 2013; Perquin et al., 2023). Methods that deal with missing data, like data imputation, assume that data is missing at random (Donders et al., 2006). However, this is typically not the case for missing values in RT series, as these reflect moments in which performance on the task was particularly poor – and as such, there are no obvious ways to deal with missing data (see Perquin et al., 2023 for a more elaborate discussion). For the current purposes, we want to avoid imputed data to cause under- or over-estimation of explained variance, and therefore excluded the respective trials.

For any missing or excluded trials, the subsequent two trials were also excluded to ensure the autocorrelations at lags 1 and 2 are only calculated on trials that directly follow each other. For the block-wise dependencies, trial indices were not altered after trial exclusion to ensure the long-term structure was preserved as much as possible. Participants were excluded from all analyses if the number of missing RTs in the full model exceeded 10% - note that this value includes both the non-responses and excluded outliers plus the two subsequent trials.

### Data analysis

#### Fitting General Linear Models

##### Full model

To quantify the total explained within-subject variance, we fit general linear models (GLMs) for each dataset on the RTs as dependent variable, using the *fitglm* function in Matlab 9 (Mathworks, 2021) with an Iterative Reweighted least squares algorhitm. All GLMs included the following predictors:

1. experimental condition(s) in the current trial, *1b. if the design included two experimental manipulations of interest: the interaction between experimental conditions in the current trial,*
2. experimental condition(s) of the previous trial, *2b. if the design included two experimental manipulations of interest: the interaction between experimental conditions in the previous trial,*
3. RT of the previous trial,
4. RT of two trials ago,
5. the linear and quadratic trends in trial number, separately for each block,
6. the linear and quadratic trends in block number,
7. the interactions between trial and block number, and
8. the interactions between trial number, block number and the experimental condition(s) – i.e., condition*trial*block including all lower-order interactions.

Points 1 and 2 represent the experimental conditions, 3 and 4 the autocorrelation, 5, 6, and 7 the long-range trends, and 8 the interactions between experimental conditions and trends (see Supplementary Materials A for the full equations).

The experimental conditions were added as categorical variables unless specified otherwise in Tables 1 and 2. Note that by using blockwise trial numbers, a new block marks a “reset” (e.g., after completing a previous block, having a break, or any other activities).

From the GLM fits, we extracted the amount of explained variance (R^2^_abc_), calculated as:

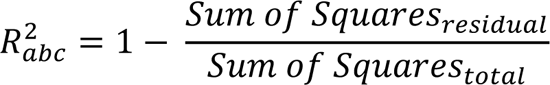

R^2^_abc_ refers to the total amount of trial-to-trial variance in RT explained by all predictors– i.e., the combination of the experimental conditions (c), autocorrelation (a), and blockwise linear trend (b) all together. Hence, the amount of unexplained variance can be calculated by 1 - R^2^_abc_.

Models that have a low number of observations, large number of to be estimated parameters, corelated predictors, or small effect sizes may lead to overfitting, which in turn leads to inflated estimates of explained variance (Yarkoni & Westfall, 2017). Therefore, we implemented a correction procedure separately for each dataset. This was done by randomly shuffling the RT series 100 times and fitting the GLMs on each of these shuffled series, and subsequently calculating the average explained variance across these 100 iterations. This procedure results in an estimate of how much each of the predictors can explain on a noise series with the same mean and variance of the RT series. If there is no overfitting at all, these estimates of explained variance should be zero. Instead, the distributions were clearly above zero (see Figure 3 – note that the estimations have been multiplied by 100 for interpretability). To correct for this bias, the mean estimate of explained variance on noise was subtracted from R^2^_abc_ for each dataset. Note that these corrected variance estimates can in princinple be negative, whith values near zero indicating that the variance explained in the actual data is comparable to an amount of variance explained by chance.

**Figure 3.**
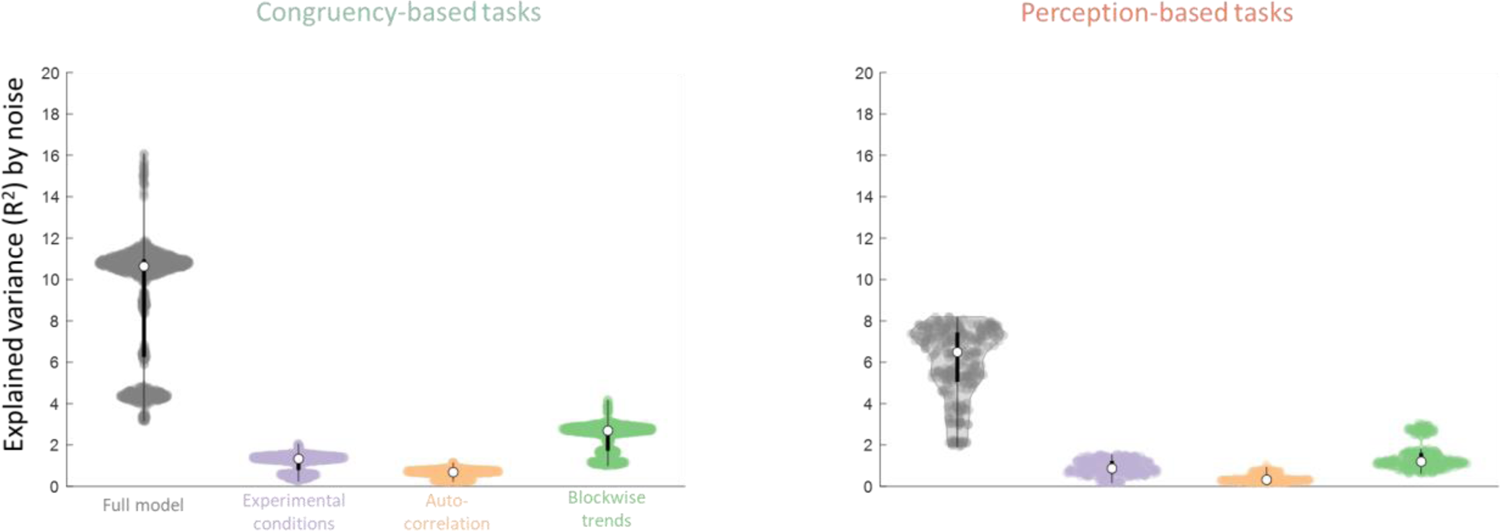
Estimates of explained variance in percentages by GLMs on shuffled RT series for the congruency-based (left) and perception-based (right) tasks. For better visibility, zoomed-in versions of the same plots are displayed on top. Each dot represents one dataset, the white circle represents the median of the distribution, and the width of the dot cloud indicates distribution density – by the three sources combined (full model or R^2^_abc_; grey) as well as separately (purple, orange, and green for experimental condition R^2^_c_, autocorrelational R^2^_a_, and blockwise trends R^2^_b_ sources respectively). Theoretically, one would expect these estimates to be zero. The distributions are clearly above zero, indicating inflation of explained variance.

##### Fitting each source of variability separately

Aside from the total amount of explained variance, we were also interested in how much variance each of the three sources – experimental conditions, autocorrelation, and blockwise trends – could explain by itself. We fitted separate GLMs by removing all other terms from the overall regression equation, and extracted explained variance for each – R^2^_c_, R^2^_a_, and R^2^_b_ respectively. These three estimates were corrected by noise in the same manner as above.

These R^2^ values reflect the variance each source explains in total while not controlling for the variance it shares with the other two sources. The variance explained by the full model (R^2^_abc_) can therefore be lower than the sum of R^2^_c_, R^2^_a_, and R^2^_b_. This is illustrated in Figure 4C: part of the explained variance may overlap between sources (i.e., overlapping parts of the circles). To quantify the amount of variance that is *uniquely* explained by each source (i.e., the size of the non-overlapping parts of the circles), we subsequently used variance partitioning analysis (Borcard, et al., 1992).

**Figure 4.**
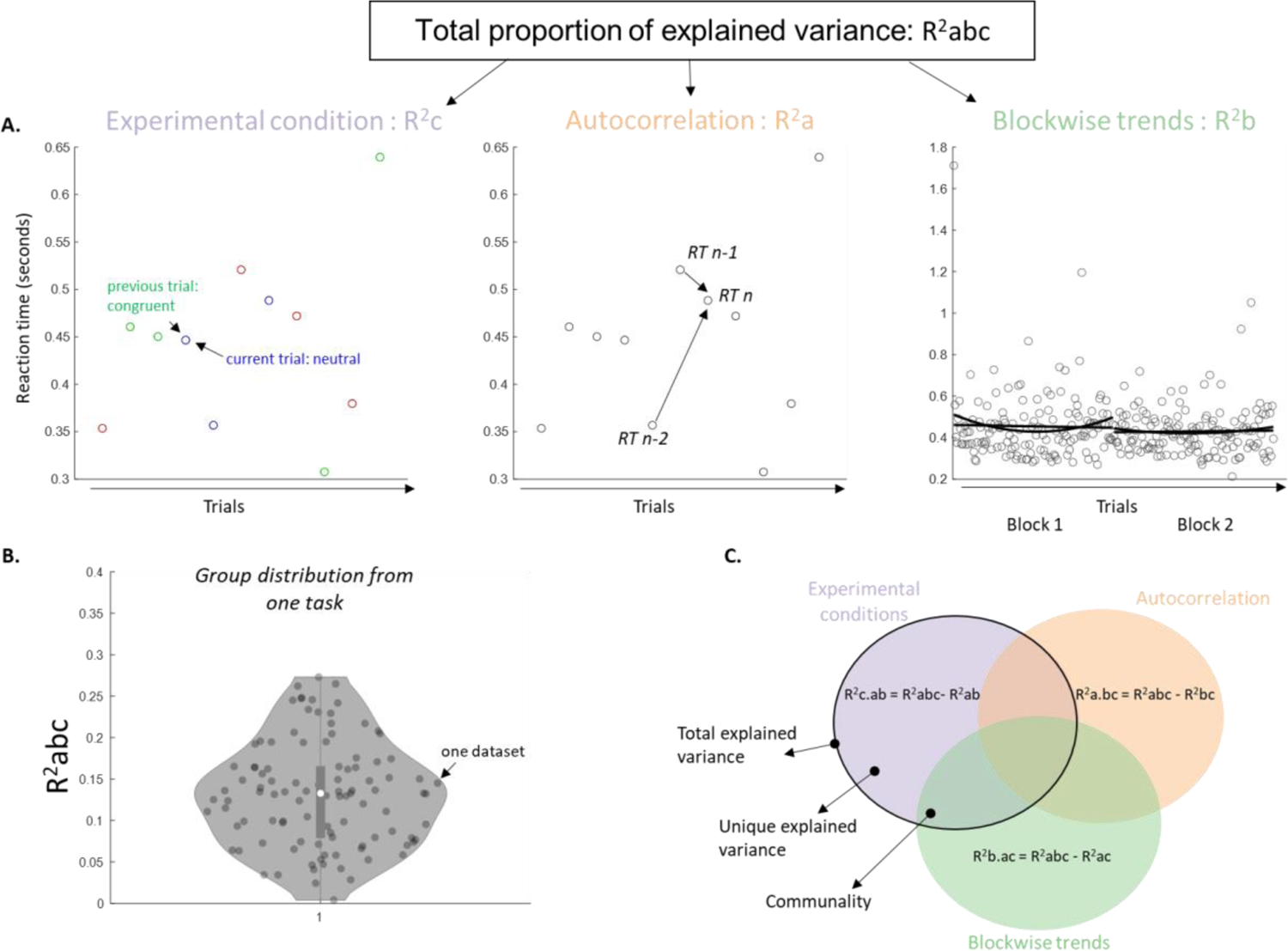
Outline of the data analysis. **A.** For each participant, GLMs were fit on RT for each of the three sources of interest: 1) experimental conditions (including the current and previous trial as predictors; left), 2) autocorrelation (including the previous and second-to-last trial; middle), and 3) linear and quadratic blockwise trends (including blockwise trial numbers as predictors; right) to extract the explained variance (R^2^) for each source. Furthermore, we fit a GLM that combined these individual predictors (“full model”) to extract the variance explained by all three sources together (R^2^_abc_). **B.** The total proportion of explained variance was extracted for each participant for each of the sources separately (R^2^_a_, R^2^_b_, R^2^_c_) as well as for all three sources combined (R^2^_abc_). Distributions of explained variance are examined on the group level, showing R^2^_abc_ as an example for one task. **C.** The unique explained variance for each source of interest is calculated by subtracting the explained variance of the other two components from the overall explained variance. The communality in turn refers to the explained variance that is shared by each combination of sources.

### Partitioning variance

#### Unique explained variance

To partition the variance, we first fitted each two-way combination of the three sources (experimental conditions plus autocorrelation, experimental conditions plus blockwise trends, and autocorrelation plus blockwise trends) and extracted their explained variance: R^2^_ac_, R^2^_bc_, and R^2^_ab_ respectively. These estimates were again corrected for noise. Next, to quantify the unique explained variance for each source, the R^2^ from the other two sources combined was subtracted from the total amount of explained variance by all three sources together. For example, we calculated the unique explained variance by the experimental conditions (R^2^_c.ab_) as:

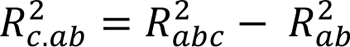

#### Communalities

Besides the unique explained variance, variance partitioning (Borcard et al., 1992) offers a quantification of the amount of shared variance (i.e., the overlapping parts of the circle; Figure 4C). For example, the communality C_ab_ between autocorrelation and blockwise trends (overlap in green and orange circle in Figure 4C) was calculated as:

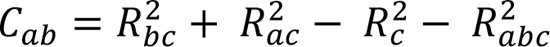

and the communalities C_abc_ between all three sources as:

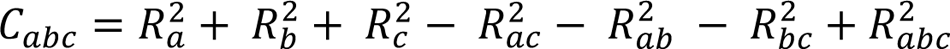

Note that theoretically communalities cannot be <0. However, for the noise-corrected variance estimates used here they can be, again with values near zero indicating no common variance.

### Individual differences

To anticipate on the results, the amount of explained variance showed large differences between individuals (see spread of data points in Figures 5). We tested whether individual differences in explained variance can be explained by differences in task performance, total RT variance, or speed-accuracy trade-offs. Three performance measures were calculated for each dataset: %-correct and mean and SD of RT on correct trials. Second, the EZ-diffusion model was used to estimate speed-accuracy trade-offs (Wagenmakers et al., 2007). This model distinguishes strategy adjustments from true performance improvements. It is based on the drift-diffusion model (DDM) by Ratcliff (1978), which aims to capture different components of decision making into different components. The EZ-diffusion model uses calculations rather than fitting procedures and is particularly suited for identifying drift rate and boundary separation effects (van Ravenzwaaij, Donkin & Vandekerchove, 2017; van Ravenzwaaij & Oberauer, 2009). It provides three key parameters: 1) drift rate (v) reflecting information processing speed, 2) boundary separation (α) reflecting the threshold to make a decision, and 3) non-decision time (Ter) reflecting perceptual and motor processes unrelated to the actual decision-making. Enhanced performance is linked to higher drift rates and shorter non-decision times, while differences in speed-accuracy trade-offs are associated with boundary separation.

**Figure 5.**
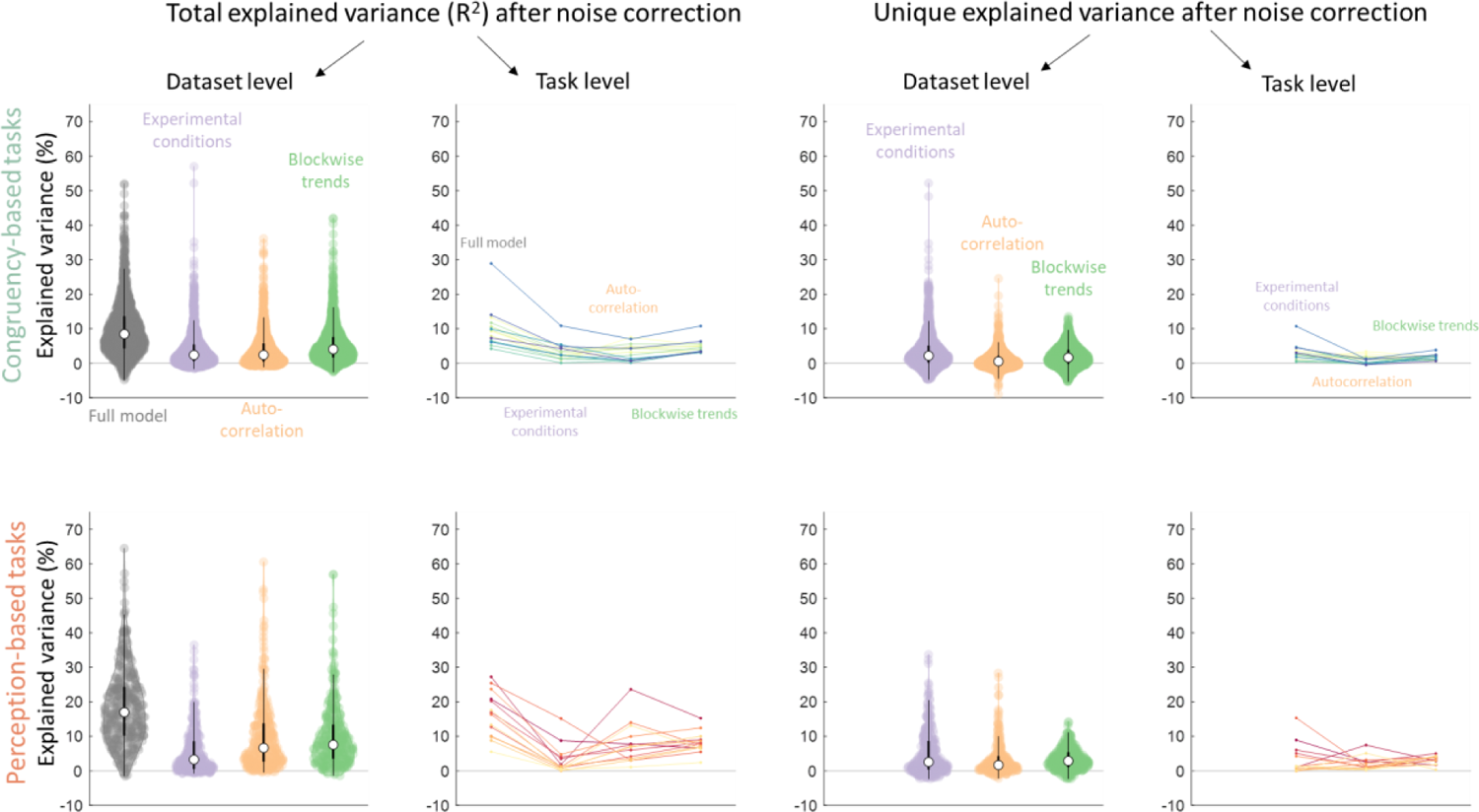
Total percentage of explained variance (left panels) and unique explained variance (right panels) in congruency-based tasks (top rows) and perception-based tasks (bottom rows) after noise correction. On the dataset level, each dot represents one dataset, the white circle represents the median of the distribution, and the width of the dot cloud indicates distribution density – by the three sources combined (full model or R^2^_abc_; grey) as well as separately (purple, orange, and green for experimental condition R^2^_c_, autocorrelational R^2^_a_, and blockwise trends R^2^_b_ sources respectively) as estimated by the dataset level GLMs. On the task level, each coloured line represents the median across datasets for each task separately across the three sources (congruency-based tasks in warm colours, and perception-based tasks in cold colours). Negative values indicate instances in which the predictors explained less variance than random noise.

Separately for each of the 30 tasks, between-subject Pearson correlation coefficients were calculated between performance (%-correct, mean and SD of RT on correct trials), drift rate, boundary separation and explained variance (total explained variance by all sources, and unique explained variance by each source separately). In total, this resulted in 24 correlation coefficients per dataset. We used Bayesian One-Sample t-tests to compare the distributions of correlations against 0. These were calculated in JASP (JASP Team, 2017), using equal prior probabilities for each model and 10,000 Monte Carlo simulation iterations. As we did not have directional expectations, t-tests were performed two-sided.

We also examined the effect of the total number of trials for each dataset. The total number of trials was correlated with explained variance (total explained variance by all sources, and unique explained variance by each source separately) across all datasets. Because the distribution of trial number was not normal, Kendall’s τ correlations were computed.

### Transparency and openness

The datasets of experiments from our labs have been made available on OSF alongside the analysis code (Perquin et al., 2023). All data from other researchers have been cited in Table 1 and 2 as well as in the Appendix. The OSF project associated with this publication contains links to these openly available datasets from other researchers, and all original papers have been cited in Tables 1 and 2 as well as Appendix A. No original research materials have been created for the current study. The study and analysis plans were not preregistered.

## Results

We analysed RT time series from a total of 30 tasks. Sixteen of them were from congruency-based tasks (1563 datasets in total, N = 1445 after exclusion; Table 2) and 14 were from perception-based tasks (350 datasets, N = 318 after exclusion; Table 3).

**Table 3.** Median percentages of total and unique explained within-subject variance plus interquartile range in brackets on the dataset level after noise correction (corresponding to the white dots in the violin plots on Figure 5). Highest values of explained variance across the three sources are displayed in bold.

|  | Total % of explained within-subject variance after noise correction | % of unique explained within-subject variance |
| --- | --- | --- |
| **Congruency-based** |  |  |
| All sources | 8.48 (9.2) |  |
| Experimental condition | 2.40 (4.7) | **2.19** (4.9) |
| Autocorrelation | 2.42 (5.1) | 0.006 (2.7) |
| Blockwise trends | **4.06** (5.8) | 1.68 (3.9) |
| **Perception-based** |  |  |
| All sources | 16.98 (14.0) |  |
| Experimental condition | 3.27 (7.9) | 2.57 (8.1) |
| Autocorrelation | 6.64 (11.0) | 1.68 (3.9) |
| Blockwise trends | **7.56** (9.7) | **2.88** (4.2) |

The main analysis centred around three sources of trial-to-trial variability, with each source consisting of multiple predictors (Figure 4A). The first source is the ‘experimental conditions’, operationalised as the condition of the current and previous trial to capture the immediate experimental effects and the sequential condition effects. Figure 4A (left panel) shows an example for one trial (left panel), in which the RT_n_ is predicted by the condition on the current trial *n* (neutral) and the previous trial (congruent). The second source is the ‘autocorrelation’, operationalised as the autocorrelation at lag 1 (= previous trial) and lag 2 (= two trials back) to capture short-range dependencies – that is, every trial *n’s RT* is predicted from previous trials *n-1* and *n-2* (middle panel). The third source is the ‘blockwise trends’, operationalised as trial-wise linear and quadratic trends across the experimental blocks to capture long-range dependency (right panel). Because our main measure is explained variance, we emphasise the numerical values as a measure of effect sizes: higher values indicate more explanatory power. No inferential statistics are used to compare the distributions. We corrected variance estimates by subtracting mean explained variance obtained from randomly shuffled data across 100 iterations per dataset (see Methods). If there were no systematic patterns in the data by either experimental conditions or temporal dependencies, the explained variance should not be higher than the explained variance explained by chance.

To calculate the proportion of explained variance, Generalised Linear Models were fit separately for each dataset, regressing RT as dependent variable. The total proportions of explained variance were extracted for each source of variance separately as well as for all three sources combined. For example, Figure 4B shows the distribution of total explained variance for the Flanker task from Hedge et al. (2018), with each dot representing one dataset and the white dot showing the median value across all datasets. Next, the unique and common proportions of explained variance were calculated for each source of variance. A visual aid for the unique and common explained variance is shown in Figure 4C in the form of Venn diagrams. For interpretation purposes, R^2^-values as well as communalities were multiplied by 100 to obtain the percentage of variance explained.

### Explained within-subject variance is low

Figure 5 shows the explained variance for all sources combined, and each source considered individually, separately for congruency-(top rows) and perception-based (bottom) tasks. Each dataset is represented by one dot in the violin (dataset level) – i.e., 1763 estimates across two types of tasks of how much trial-to-trial variance is explained by the three sources. The density of the violin shows where most datasets are located, and the white dot represents the median value across all datasets. The numerical values of these medians are provided in Table 3. The total proportion of variance (left panels) explained by the experimental conditions was less than a third of the total explained variance; in other words, temporal dependencies explained more variance than experimental conditions. Moreover, the total contribution of experimental conditions was <10% for 1313 out of all 1445 analysed RT series (91%) in the congruency-based tasks, and for 255 out of 318 RT series (80%) in the perception-based tasks. Numerically, blockwise trends explained the highest proportion of variance. The unique variance (right panels) explained by autocorrelation and blockwise trends were numerically substantially lower compared to their total explained variances (see Table 3). This reduction was only minimal for the experimental conditions. Numerically, median explained variance was higher for the perception-compared to the congruency-based tasks.

It may be possible that there are differences between the different tasks that would skew the dataset level distributions. We therefore we calculated the group medians separately for each task to examine consistency across tasks. These are shown in the task level panels in Figure 5, with the medians for each task being represented by one line. Violin plots across participants separately for each task can be found in Supplementary Materials B. On the task level, the three sources collectively explained on average 1/7 to 1/10 of the total trial-to-trial variance. Thus, the largest part of the variance remained unexplained when summarising the results across tasks.

### Short- and long-term dependency, but not condition, share explained variance

We also extracted the amount of explained within-subject variance that was shared between sources (i.e., communality; overlapping parts of the circles in Figure 2C). There was virtually no overlap between the variance explained by the experimental conditions and the temporal dependencies (see Figure 6 for distributions). For example, the variance commonly explained by experimental conditions and autocorrelation (blue) was on average .02 % of the total within-subject variance. The other communalities involving experimental condition (grey and pink) were also negligible, both in the congruency- and perception-based tasks. The median communality value between autocorrelation and blockwise trends was 3.98% and 1.92% for congruency- and perception-based tasks, respectively. This indicates that shared explained variance between short- and long-term dependencies amounts to about 2-4% in the total trial-to-trial variance. Overall, explained variance originating from experimental conditions was thus largely unique, whereas the two temporal sources shared explained variance.

**Figure 6.**
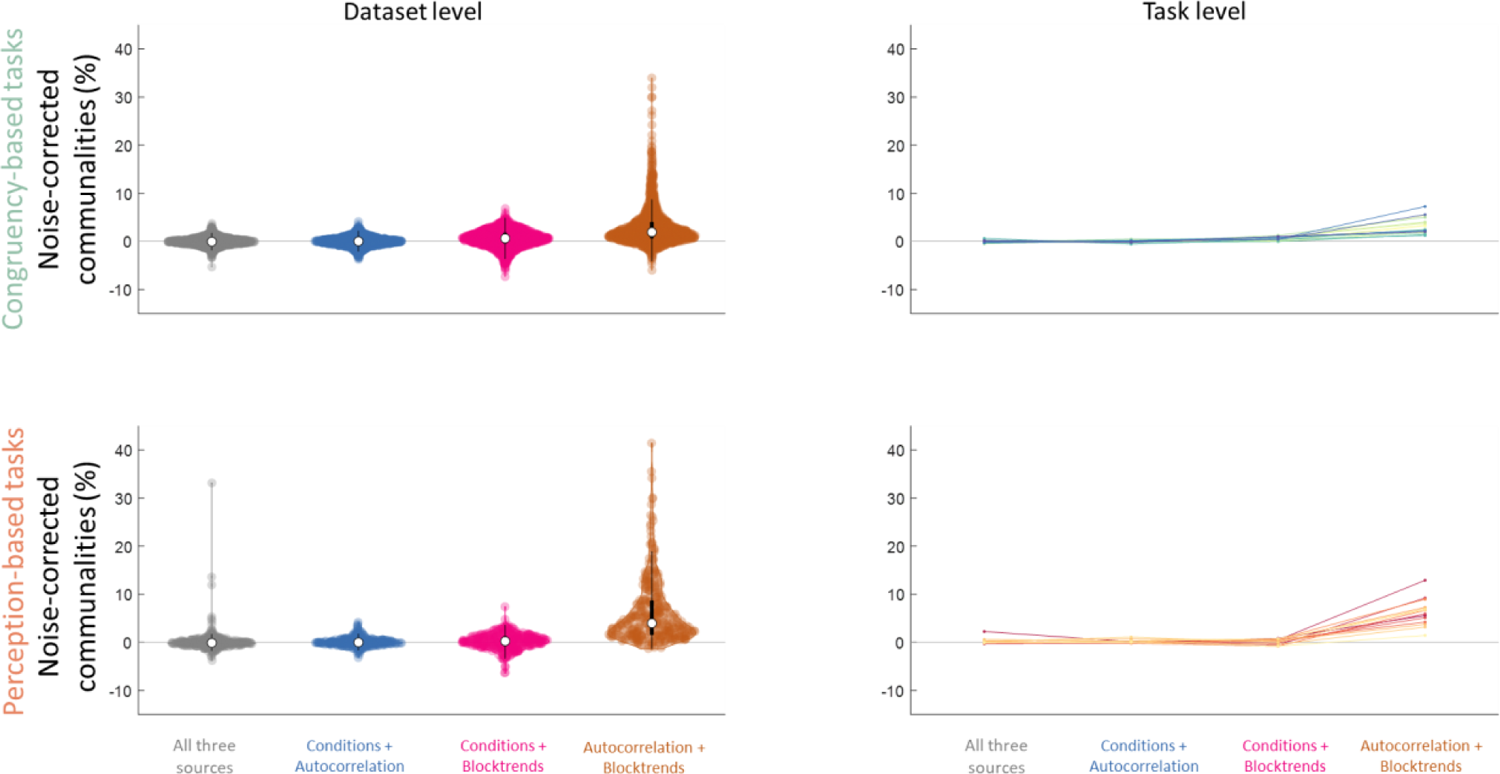
Percentage of common explained within-subject variance by different sources: experimental conditions plus autocorrelational (blue), experimental conditions plus blockwise trends (pink), autocorrelation plus blockwise trends (brown), and all three combined (grey), shown for the congruency-based (top) and the perception-based (bottom) tasks. The communalities are shown both on the dataset level (left, with each dot representing one RT series) and task level (right, with each coloured line representing one task). The communalities between autocorrelation and blockwise trends are above zero, indicating an overlap of explained variance. The other three distributions are near zero, indicating negligible overlap. Communalities were calculated on the noise-corrected estimates of explained variance.

### Effect of accuracy

RT distributions of correct and incorrect trials are known to differ both in mean and variance. We therefore ran a model to estimate the contribution of accuracy to trial-to-trial variability (Figure 7). Explained variance was increased with 1.1% by for the congruency-based tasks and with 1.8% by the perception-based tasks (dark purple violins). The increase in explained variance differed noticeably across tasks – especially in the congruency-based tasks (right panels).

**Figure 7.**
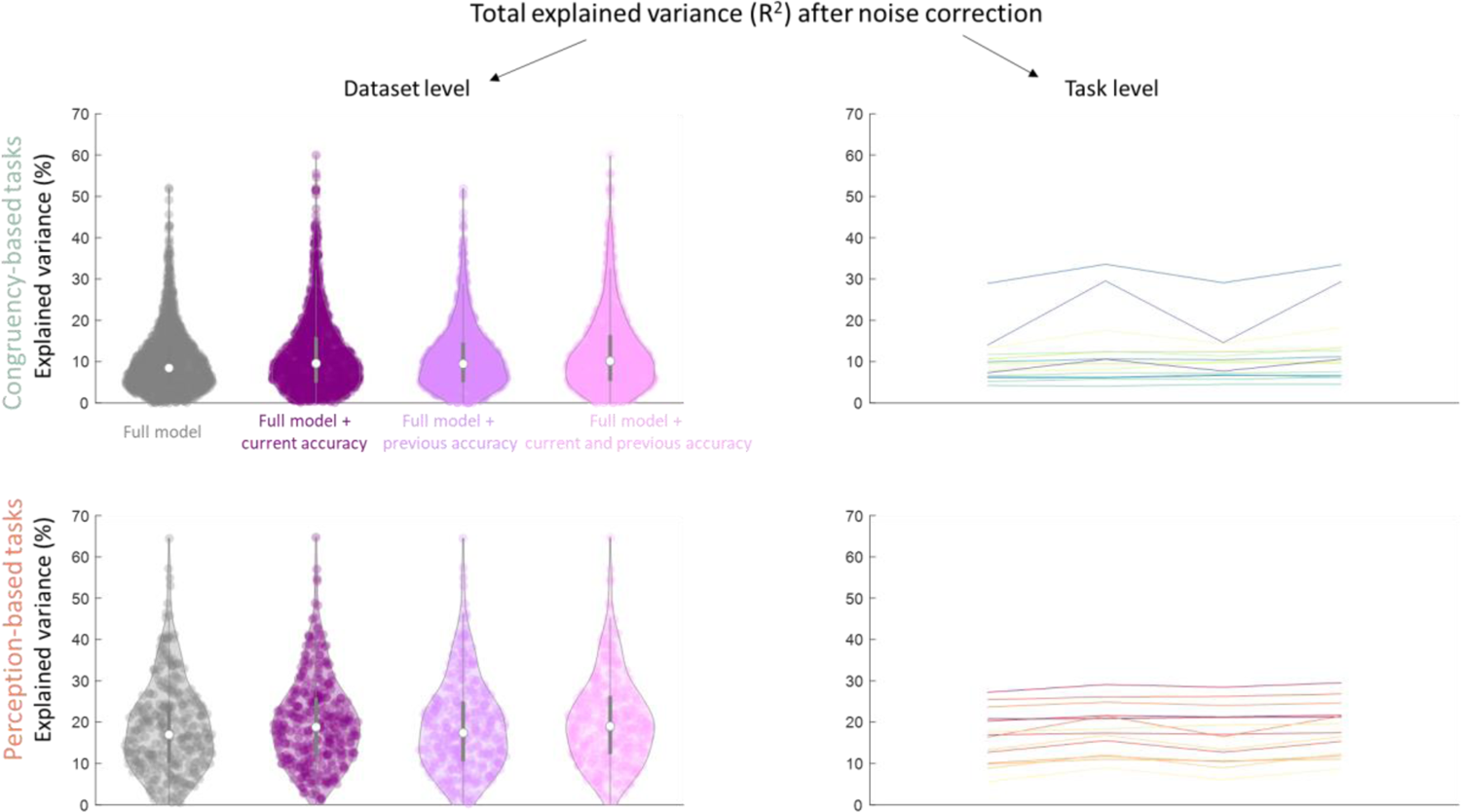
Percentage of noise-corrected within-subject variance by the full model (grey, also plotted in Figure 5), by the full model plus accuracy on the current trial (dark purple), by the full model plus accuracy on the previous trial (lavender), and by the full model plus accuracy on the current and previous trial (soft pink). The explained variances are shown both on the dataset level (left, with each dot representing one RT series) and task level (right, with each coloured line representing one task).

Next, as it is known that accuracy on the previous trial can influence RT, we ran an additional model that included accuracy on the previous trial (lavender violins). Compared to the full model, this increased the explained variance by 1.0% for the congruency-based and .5% for the perception-based tasks. Finally, a model including both current and previous accuracy (pink violins) explained 1.6% and 2.0% more variance compared to the full model respectively for congruency- and perception-based tasks compared.

### Control analyses confirm that results are robust

We conducted several further control analyses to inquire whether the above results are robust to the choices made during data analysis. First, we assessed the influence of different outlier exclusion criteria and commonly applied transformations of RT. Second, we verified that task-irrelevant exogenous elements in the design (e.g., basic stimulus characteristics or inter-trial interval durations) contribute very little to the overall RT variance. Thirdly, the main analyses were rerun after excluding datasets for which the predictors showed signs of severe multicollinearity (marked with an asterisk in Table 2). None of these aspects affected the conclusions we have presented: their numerical influences were small. Finally, we checked if the main results change when removing the variance of experimental conditions and blockwise trends from the autocorrelations – by first regressing RT on experimental conditions and blockwise trends, and subsequently using the residuals as predictors for the previous RTs. This procedure reduced the total explained variance by the autocorrelation, but not the unique explained variance nor the explained variance by all predictors combined – confirming the results of the communalities analysis. We report all control analyses in the Supplementary Materials C.

### Mixed models

There has been an increasing interest in linear mixed models (LMMs) to study the effects of experimental conditions (Singmann & Kellen, 2019). These can include both fixed effects (which remain constant) and random effects (which signify variance across the levels of grouping variables in the dataset). In typical between-subject approaches, they can therefore explain variance between participants which GLMs will assign as error variance. When one wants to take advantage of this feature of LMMs, all participants must thus be included in one overarching model, so it can assign as much of the variability between participants as possible to the random factors. Note that in contrast, our GLM approach used subject-wise GLMs, thus fitting each participant’s data independent of all others and accordingly allowing each GLM maximal flexibility to fit individual data. Therefore, employing an LMM to our multi-study/multi-participant datasets (see Figure 3) has a different focus than the analysis we have presented. We present LMM analyses for the current data in the Supplementary Materials D. The LMM explained overall more variance than the GLM approach, but still left most of the total variance unexplained. This difference was due to higher amounts of explained variance related to short- and long-term dependencies. In contrast, experimental manipulations explained less variance in the LMM than in the GLMs. We discuss potential pitfalls in comparing these numerical comparisons between the two model approaches in section *Model form* in the Discussion.

### Individual differences in performance have an unsystematic and small impact

The amount of explained variance showed large differences between individuals. One might speculate that these are driven by performance – e.g., a well-performing participant might be less influenced by temporal effects and show larger experimental effects – or by the variability itself – e.g., if a participant is more variable, there is also more potential variability to explain. To test whether individual differences in performance or speed-accuracy trade-offs might explain differences in explained variance, Pearson correlation coefficients were calculated separately per task between five measures of performance/speed-accuracy (total %-correct, mean RT on the correct trials, SD of RT on the correct trials, and drift rate and boundary separation as calculated by the EZ drift diffusion model) and four measures of explained variance (total by the full model, and uniquely explained by each of the three sources). If individual differences in speed-accuracy trade-offs can (partially) explain individual differences in explained variance, this would be reflected the distributions of coefficients being different from zero.

Figure 8A shows the 20 distributions along with the Bayes Factor (BF_10_) on top. Five of the 20 distributions showed clear Bayesian evidence in favour of a positive correlations between measures of performance and explained variance, seven distributions showed clear Bayesian evidence against correlations, and eight remained indeterminate. The largest correlation was found between drift rate and explained variance by condition (median r = .23). This means that on average 5.3% of the between-subject variance in explained variance by condition can be attributed to individual differences in drift rate – with higher drift rates indicating more explained variance. Explained variance for the other four distributions ranged from .6 to 2.3%. We conclude that performance might have a small impact on explained variance, though there is no clear consistency across the directions of effects.

**Figure 8.**
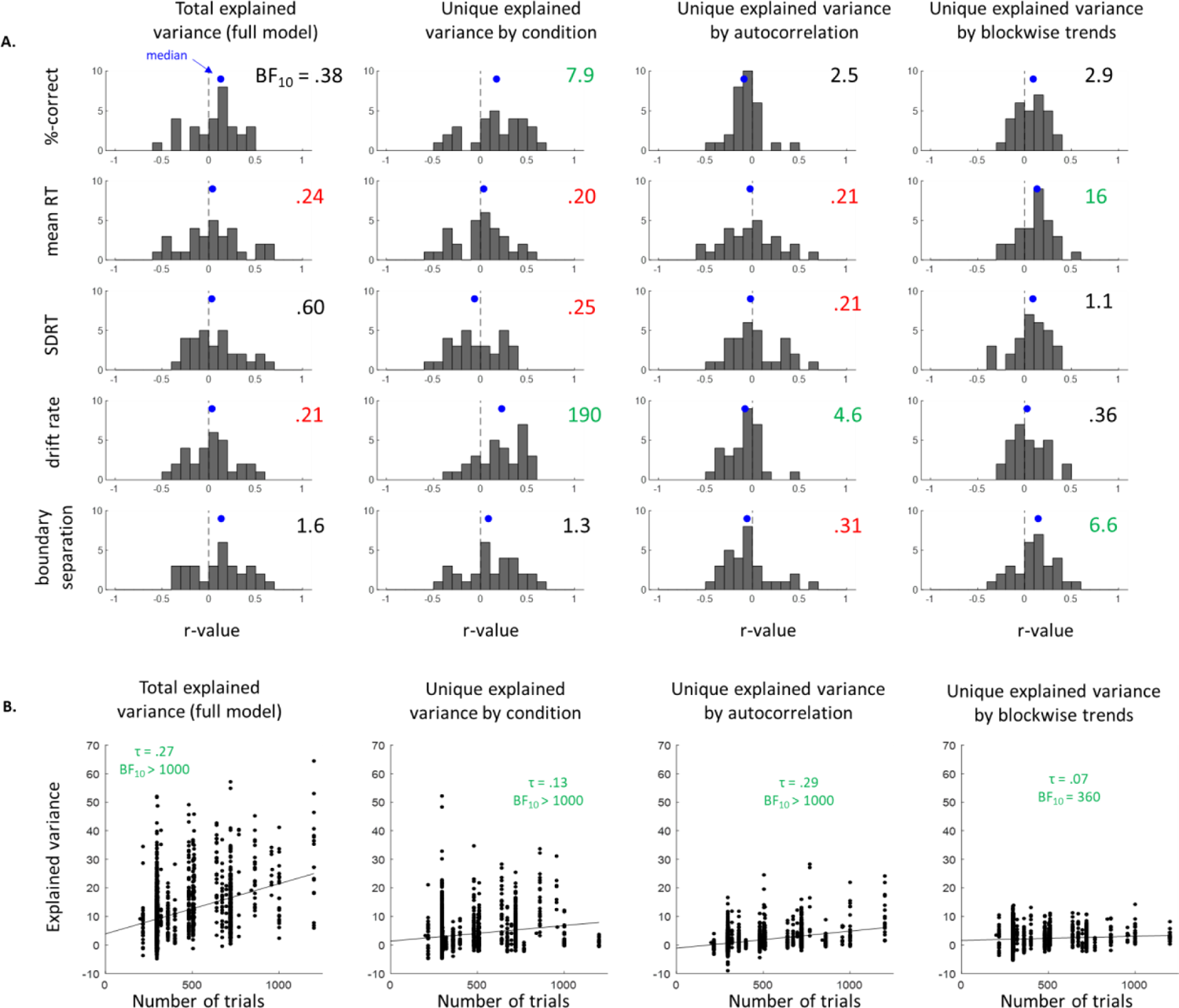
**A.** Distributions of Pearson correlation coefficients across tasks, across four measures of noise-corrected explained variance (columns) and five measures of performance and speed-accuracy trade-off. Each distribution was tested against zero (dotted line). The corresponding Bayes Factors are shown on top. Bayesian evidence favouring the presence of a correlation between explained variance and performance are shown in green (BF_10_ > 3), evidence favouring the absence of a correlation (null effect) are shown in red (BF_10_ < 1/3), and indeterminate Bayes Factors are shown in black. The median r-value of each distribution is represented by the blue dot. **B.** Bayesian correlation analyses between the number trials and four measures of noise-corrected explained variance. The τ-value and BF_10_ are shown on top.

We also tested the relationship between total trial number and explained variance, by correlating the same four measures of explained variance with total trial number across all datasets (Figure 8B). These correlations were systematically present and positive, meaning that both variance explained by experimental conditions as well as endogenous temporal dependencies were stronger with increasing trial numbers. However, the explained variance was likewise small (ranging from .5 to 7.3%).

## Discussion

We systematically quantified the variance in trial-to-trial RTs explained linearly and additively by experimental conditions and short- and long-range temporal dependencies systematically across a sample of congruency- and perception-based tasks. We focused on studies that are representative of their respective fields, as they are specifically designed to study sensory and cognitive phenomena, and do not aim to increase or reduce RT variability. Specifically, GLMs were conducted as a means to test how much variance may be explained by conventional models in cognitive neuroscience. Although the experimental manipulations in these tasks typically induce robust differences between conditions on the level of averaged single subject and group data, they did not explain more variance than temporal dependencies such as autocorrelation and blockwise trends, which are traditionally considered nuisance. In addition, there was no overlap of the variance explained by experimental conditions and temporal dependencies. Proportions of explained variance were similar across the two task sets. Hence, while RTs may change systematically over time, RT differences between conditions are stable. Perhaps most strikingly, our results show that GLMs leave the largest part of within-subject variability unexplained by the sources analysed in the present work.

We analysed data from two types of tasks that are often used in experimental psychology and cognitive neuroscience: congruency-based tasks and psychophysical perception tasks. Some have argued that psychophysical tasks are particularly suited for inducing robust effects across participants (Smith & Little, 2018), given the frequent use of individually calibrated stimuli (which enhances the condition effect at the individual level) and the collection of small but well-trained samples (which reduces the error variance compared to testing naïve participants). In contrast, most congruency-based tasks employ fixed group designs and rely on larger, untrained participant samples. These differences are also reflected in the datasets analysed here: congruency-based tasks included more participants, while perception-based tasks featured more trials and blocks. In total, seven studies used individually calibrated stimuli, but their variance patterns were not different from other tasks in any obvious way (see Table 3). However, we do not have the means to systematically compare the two types of tasks, nor any differences between tasks in general, and only conclude that the overall conclusions hold up across both.

### Experimental effects versus intra-individual variability

Typically, significant differences between experimental conditions are interpreted in the form of *‘responses are faster in condition A than in condition B’*. Such interpretations emphasise the *direction* (not the magnitude; also see Rouder & Haaf, 2021) of the difference between condition-wise *summary statistics (such as means and medians)* that are usually further *summarised across participants*. In this form, many experimental effects, including those analysed in the current study, are considered to be highly robust even if the magnitude of a condition-difference may be small (Rouder et al., 2019; Rouder & Haaf, 2018). As such, these tasks are a popular means to test group distributions between conditions, though the amount of explained variance (or magnitude of effect) often remains unreported. The group distributions are typically analysed under the assumption that the effects of interests are linear and additive (Blanca et al., 2018).

Yet our results show that even if the group-wise effect of an experimental manipulation is replicable, the moment-to-moment variability of an individual participant’s responses remains largely unpredictable with the same linear additive models. For example, a well-known cognitive congruency effect (the Stroop effect) was replicated in each of 20 independent participant pools in the cross-laboratory experiment “Many Labs 3” (Ebersole et al., 2016), and it has even been said that “Everybody Stroops” (Haaf & Rouder, 2017). If we were to test a naïve participant on this task and predicted that their average behaviour will be faster in the congruent compared to the incongruent condition, our prediction would almost certainly be correct. However, as shown by the present analysis, our ability to predict the RTs of individual trials remains poor even for this particular task. Hence, the robust effects of experimental conditions in typical neurocognitive tasks do not immediately translate to explanatory power for RT at the single-trial level.

We also examined the increases in explained variance when adding the current trial, as distributions of correct and incorrect RTs are known to differ (Ratcliff, 1978; Bolsinova & Molenaar, 2018). The increases were modest, although there were stark differences across tasks. These results should be interpreted with caution though. It is known that experimental conditions can cause changes in RT, as they can be manipulated directly by an experimenter. It is also likely that time can cause changes in RT, as their correlations cannot be explained bi-directionally (i.e., RT cannot cause changes in time). Accuracy itself is a measure of within-subject variability, so one cannot say that accuracy causes differences in RT – that would mix up the explanans and explanandum. In contrast, accuracy on the previous trial can be a possible source of within-subject variability: participants slow down after they realise they made an incorrect response (post-error slowing; Rabbitt, 1966). The increase in explained variance by previous trial was modest. However, post-error slowing is highly task-depended, as it requires either awareness of the error or explicit feedback. Most of the perception-based tasks specifically do not meet this requirement.

### Temporal dependency in variability

Short- and long-range temporal dependencies explained more variance than the experimental conditions. Still their total contribution to explained variance low, implying that knowledge of the precise temporal patterns of variability (e.g., the strength of autocorrelation) provides only poor prediction of the individual RTs.

For practical reasons we limited the analysis on specific (first- and second-order) forms of temporal structures, though some studies have described higher-order structures in RTs. For example, one line of research has described long-range or scale-free patterns in behaviour (e.g., Gilden, 2001; Kello et al, 2007; Thornton & Gilden, 2005; Van Orden et al., 2003). Often, such structure has been quantified using frequency analysis (Box et al., 2016) and some studies have claimed that long-range structures explain “a fair fraction” of the total variance, and in some cases more than the experimental conditions (Gilden, 2001, p. 54). These analysis methods come with various controversies, and the explained variance cannot be attributed to defined timescales, which limits its interpretability (see Wagenmakers et al., 2004 for a comprehensive overview; also see Farrell et al., 2006). In addition, entering such structures into an analysis of shared and unique variances is not straightforward. As such, it remains open whether higher-order temporal structures explain more variance than the autocorrelations and block-wise trends investigated here.

### Generalisability of findings

For our study, we analysed data from studies on cognitive control and sensory perception, using tasks that are well representative of their respective research fields. Proportions of explained variance were highly similar across congruency- and perception-based tasks. Whether our findings generalise to other types of tasks and disciplines remains an empirical question. However, congruency- and perception-based paradigms tend to produce condition differences in RT on the group level, and both typically include relatively high trial numbers to minimise measurement error compared to other fields (e.g., motor control, memory), and are often associated with error rates that are either low or fixed at the group level (e.g., by using staircase procedures). It seems unlikely we would find larger proportions of explained variance by experimental manipulations in tasks that do not share these characteristics. Furthermore, the potential reasons for why our behaviour may be variable seem largely universal (see section *Other predictors of trial-to-trial fluctuations* below). Overall, the low proportion of explained variance may thus be a general feature of neurocognitive tasks. Similar conclusions have been drawn in animal work (Musall et al., 2019). This may reflect that the ongoing experimental task is only a small part of the plethora of activity the brain is processing at each moment in time, even in highly controlled laboratory environments.

The present analyses only include tasks that involve manual button presses. RT produced by other modalities are quantitatively different from button presses – e.g., RT latencies from button presses are longer than those from saccadic eye movements). More importantly, they can be *qualitatively* different. For example, in rapid action selection tasks, manual RT had longer and more variable non-decision times – reflecting sensory processing and motor output – compared to saccadic RT (Bompas et al., 2017). If different types of responses are qualitatively different, we cannot easily speculate if the specific proportions of explained variance found in the current study would generalise to trial-to-trial RT from other output modalities. Still, saccadic responses are also known to be highly variable (Carpenter, 1999; Noorani & Carpenter, 2016), and it may be fair to assume that large proportions of this variability would be unrelated to the task.

It is possible that temporal predictability is higher in sensorimotor actions other than the ones we scrutinised in the present study, particularly in tasks without experimental conditions such as tapping to a metronome (see Perquin et al., 2023 for an overview of autocorrelations in RT series from various tasks with and without experimental conditions). However, even in these tasks without experimental conditions, the largest proportion of the RT variance remained unexplained – meaning the trial-to-trial responses were still largely unpredictable.

By combining datasets from different experiments, the present approach allows for large-scale replication; dataset can be considered as another replication to the same research question. However, not all are *independent* replications, because several datasets were acquired from the same participants who performed multiple tasks in one study (Figure 3). The potential bias of including participants multiple times is likely low, because it is known that neither effects of experimental manipulations (Hedge et al., 2018; Rey-Mermet et al., 2018) nor temporal dependencies (Perquin et al., 2023) generalise across tasks – i.e., knowing the congruency effect or autocorrelation of one participant in a Stroop task tells us nothing about their effect or autocorrelation in the Flanker task. Still, there is an aspect of hierarchy to the included datasets that our approach does not take into account. To our knowledge, there are currently no methods that quantify variance solely at the level of a single participant while taking group-level aspects (e.g., task, duration, paradigm type) into account. Instead, we used LMMs to get a common estimate of explained variance across datasets, as well as one estimate of explained variance for each task (reported in Supplementary Materials D). With the large numbers of predictors in our present analysis, fitting LMMs is often impossible, because the high number of estimated parameters and their dependencies cannot be estimated even from the huge amounts of data in our study, resulting in convergence errors during model fitting. Furthermore, the use of standardised effect sizes for LMMs is controversial, and there is no obvious equivalent to R^2^ (Rights & Sterba, 2019). The LMM results should thus be taken with caution.

### Explaining RT variability

So far, we have emphasised that the analyses implemented here leave about 80-90% of the variance unexplained. The obvious question that arises hence is: what does explain the remaining variance? First, it is possible that the current sources explain large parts of RT variability, but only in non-linear structures. Second, it is possible that fluctuations in RT largely arise from fluctuations in preceding neurobiological processes not measured in the current datasets. Third, it is possible that RT variability is inherently random. These are not necessarily incompatible with each other; they might all be true to some extent. The current research does not provide empirical evidence in favour of or against any of these three explanations. We discuss all three below.

### Explanation 1: we need better models

One of the larger limitations of the current study is that the reported results are only as valid as our model choice. All the results and conclusions presented in the current study are based on general linear models. Such models are the bread and butter of cognitive neuroscience (Blanca et al., 2018), and our results highlight their limitations in single-trial prediction. Historically, linear models have been perceived as more interpretable, biologically plausible, and better suited at testing the types of hypotheses common in neuroscience than non-linear models (but see Ivanova et al., 2018 for a rebuttal of this perception).

The choice of GLM makes the current results comparable across the three different sources as well as across the different datasets. However, it is certainly possible that the general linear model is poor at explaining single-trial variance, because it cannot capture non-linear and non-additive trends that might be present in the trial-to-trial RT series. Thus, our analysis approach may have missed systematic trial level variations attributable to the experimental conditions and temporal dependencies. This is an important limitation as it is unclear how much of the possible relationship between sources and RT fluctuations are linear, and how much are non-linear. Background knowledge is lacking here, as behavioural data has modelled almost exclusively with linear models. As such, there is an almost infinite number of non-linear approaches to pick from but a lack of a priori knowledge to select the most appropriate ones.

Approaches that may identify non-additive effects, like generalized additive models (GAMs), or non-linear effects, like support vector machines (SVM), have seen increasing usage in neuroscience (e.g., Groot et al., 2017; Jin et al., 2019; 2020). On a practical level, such approaches are unfeasible in the current study as they commonly require regularisation parameters separately for each dataset that need cyclical cross-validation and re-fitting until the best fit is found. Consequently, we cannot rule out the possibility that the three sources do explain large parts of the variance in structures not captured by the GLM. If this is indeed the case, one might also wonder if non-linear and non-additive models may have a more important role in group analyses depending on the research goal (Ivanova et al., 2018). Exploration of non-linear and non-additive model approaches may explain variance that is not captured by conventional linear models, and reveal nuances in relationships between variables (e.g., Bolsinova & Molenaar, 2018, showing that RT and accuracy are related to each other in non-linear fashion).

### Explanation 2: we need to measure endogenous processes

Of course, there are other potential sources of trial-to-trial fluctuations that we have not analysed – with the most obvious being of endogenous nature. In the following we discuss two research lines that have investigated potential endogenous contributions to fluctuations in behaviour.

#### Predicting behavioural fluctuations from attentional state

One line of research explains fluctuations in performance as a result of changes in attentional state – e.g., “I was slow to respond to that other car *because* I was not paying attention”. If this is view was correct, attentional states would be one avenue for predicting upcoming responses (e.g., Baldwin et al., 2017). On the one hand, attention may refer to a global state, relating to states as arousal or motivation. Indeed, when participants report to be attentionally off-task (e.g., mind wandering, drowsiness), their average RT variability in trials just before this report is higher compared to when they report to be on-task (Anderson et al., 2021; Laflamme et al., 2018; Seli et al., 2013; Thomson et al., 2014). However, effect sizes tend to be small: fluctuations in attentional state reports can typically only explain ∼2-7% of the total trial-to-trial RT variability – though these reports appear to explain more variance in accuracy (Kane et al., 2021). Furthermore, subjective reports are typically measured retrospectively (e.g., “what was your attentional state just before this question appeared?”), but *upcoming* trial-to-trial variability as well as slow and incorrect responses seem to be consciously inaccessible (Perquin et al., 2020), meaning that metacognitive states likely have little to no predictive power.

On the other hand, attention may refer to the more specific mechanism of spatial orienting, which can be experimentally manipulated by introducing attentional cues and increasing predictability of the stimulus – though note that: spatial attention can also vary from trial to trial due to changes in arousal (see Renart & Machens, 2014; Roelfsema 2004). Attentional cues can decrease variability in behaviour and brain activity (Azari et al., 2009; Cohen & Maunsell, 2009; 2010; Ledberg et al., 2012). Likewise though, this effect appears likewise stronger on accuracy than on RT – for example, Azari et al. (2009) found that valid cues increased accuracy compared to invalid cues, but did not significantly affect RT. Overall, while attentional state is consistently associated with behavioural variability, it is unlikely to explain large proportions of the trial-to-trial fluctuations in RT.

#### Predicting behavioural fluctuations from neuroimaging data

Variability is present throughout the nervous system from phenomena such as synaptic transmission to cross-network connectivity (for comprehensive reviews, see Ermentrout et al., 2008; Faisal et al., 2008; Garret et al., 2013; Renart & Machens, 2014; Waschke et al., 2021), and is largely independent from the task at hand (Musall et al., 2019). Variability in neural activity prior to registration of a stimulus has been associated with fluctuations in subsequent actions in response to that stimulus across human and animal studies (e.g., Cohen & Maunsell, 2009; Everling et al., 1997; Parto Dezfouli et al., 2018; Supèr et al., 2003; Womelsdorf et al., 2005; Zhang et al., 2008). Correlations between fluctuations in behaviour and brain state have been found across multiple time scales, from ultra-slow (Monto et al., 2008; Weissman et al., 2006) to alpha, beta, and gamma fluctuations (e.g., Bompas et al., 2015; Gonzalez Andino et al., 2005; Hamm et al., 2010; Perfetti et al., 2011). These fluctuations may influence the encoding of sensory stimuli, as evidenced by electroencephalogram (Iemi et al., 2019) and voltage imaging data (Davis et al., 2020; Liu et al., 2022). Oscillatory phase may also play a role (e.g., Busch et al., 2009; Schroeder & Lakatos, 2009; VanRullen, 2016; but for negative findings, see Benwell et al., 2017; Bompas et al., 2015: Ruzzoli et al., 2019). Functional connectivity within and between task, sensory, and default networks also contributes to behavioural variability (e.g., Thompson et al., 2013; Zuberer et al., 2021). Furthermore, fluctuations in behaviour have been associated with other biological processes such as heartbeat (e.g., Park et al., 2014; Salomon et al., 2016), respiration (e.g., Johannknecht & Kayser, 2022), and stomach activity (e.g., Rebollo & Tallon-Baudry, 2022; Richter et al., 2017).

Collectively, this makes for a very heterogeneous body of work relying on diverse paradigms, analysis methods, and time scale. However, most studies have not reported which proportion of the total variance is explained by the respective source of interest, and endogenous and exogenous sources of variability are commonly conflated. Studies that do report proportions of explained variance point to low effect sizes: 1-4% of endogenous saccadic RT variability could be explained by oscillatory power across the task-network (Bompas et al., 2015), while total cortical transmission times across task-networks could explain on average 8% of simple RT variability (Paraskevopoulou et al., 2021). It is thus unlikely that there is either a single process or a small number of processes that will explain a large proportion of behavioural variability.

### Explanation 3: we cannot explain variability

It is possible that part of moment-to-moment variability is inherently random, and thus cannot be explained from any other systematic source. By definition, such random fluctuations would be truly unpredictable.

The bar of empirical evidence for this explanation is much higher, as it would require examining all possible sources and all possible source-behaviour relationships. Still, possible benefits of randomness have been highlighted previously in the literature. Random variability has been highlighted as a possible means to facilitate creative, flexible, and strategic behaviour and even free will, as randomness in behaviour prevents stereotypy and monotony (Carpenter, 1999; Renart & Machens, 2014). Unpredictable variability might be advantageous in contexts in which perpetuation is unwarranted, be it fleeing from a predator, exploring the environment, deciding between similar options, motor learning, or curiosity (e.g., Briffa, 2013; Chang et al., 2017; Dinstein et al., 2013; Shahan & Chase, 2002; Sternad, 2018).

### Perspectives for future research

A comprehensive understanding of human behaviour as such requires an understanding of the contributing endogenous fluctuations (see Evans et al., 2020; Uddin, 2020). Several approaches may be beneficial for addressing this question.

First, endogenous and exogenous variability are often not fully separated; any time-varying task-element (e.g., interleaved conditions, inter-trial intervals, breaks) can potentially affect behaviour. When the interest lies on endogenous – i.e., spontaneous – fluctuations, it is therefore important to minimise exogenous elements. This can for example be achieved by using highly repetitive tasks, such as tapping to a fixed tone or rapid action selection tasks that include easily perceivable stimuli and simple actions. When exogenous factors are still present, they will first need to be removed from the trial-to-trial fluctuations prior to data analysis (e.g., Benwell et al., 2018; Bompas et al., 2015).

Furthermore, multi-modal measurements that concurrently assess neural and bodily signals will be central to identifying which of the many possible neural and bodily processes uniquely contribute to behavioural fluctuations. An important aspect for the identification of variance sources will be that studies routinely report the proportion that each source explains of the total variance on the single trial level. This requires statistical approaches that focus on single trial data rather than summary statistics, and that analyse RT as a continuous rather than categorical (e.g., median split) variable. Such approaches may also benefit from non-linear models (Ivanova et al., 2018).

Both behavioural variability such as the mean and variance across an entire session (Hedge et al., 2018; Hultsch et al., 2002; Hultsch et al., 2000; Perquin & Bompas, 2019; Perquin et al., 2022; Saville et al., 2011; 2012) as well as temporal dependencies in entire RT series (Perquin et al., 2023) exhibit considerable stability across time and are thus trait-like. This makes them well-suited for research about individual differences. Furthermore, individual differences in variance structure may be a reliable trait – that is, people reliably differ in how consistent they are on congruent versus incongruent trials in cognitive tasks (Williams et al., 2022). Various neuropsychiatric diseases have already been associated with increased variability compared to healthy populations, such as ADHD (Kofler et al., 2013; Tamm et al., 2012), Alzheimer’s Disease (Tales et al., 2012; Tse et al., 2010), as well as schizophrenia, depression, and borderline disorder (Kaiser et al., 2008; also see Dinstein et al., 2013). Furthermore, recent work has suggested that neural activity associated with experimental factors is highly consistent across participants, while neural activity associated with fluctuations in task performance has high inter-subject variability but is stable within individuals over time (Nakuci et al., 2022). These findings suggest that neural fluctuations relating to performance could in itself be an individual trait, though their reliability over time and contexts remains an open question.

More complex computational models capture variability in latent variables. For example, accumulation models (e.g., Bogacz et al., 2006; Ratcliff et al., 2016) identify a drift rate of evidence accumulation for the ongoing decision and a threshold at which decisions are committed to. These types of models have been successful at simulating RT distributions and capturing the relationship between RT and accuracy and may even be restricted with neuroimaging data (e.g., Turner et al., 2013; 2016). However, their contribution to understanding trial-to-trial fluctuations is limited, because a) they include noise parameters agnostically fitted solely for the purpose of computational tractability, b) they posit, by definition, that RT series are random over time, and c) they cannot, for this reason, predict a single trial’s random walk or evidence accumulation (also see Evans et al., 2020; Evans & Wagenmakers, 2019). One avenue to understanding trial-to-trial fluctuations might be to expand such models, for example, by incorporating temporal dependencies and implementing single-trial estimates of latent variables. We note that accumulation models may implement within-trial (drift rate) and between-trial (e.g., between-trial standard deviation of drift rate) variability, representing fast fluctuations in accumulation within single trials and slower fluctuations in global processes across trials respectively. However, our current analyses do not allow for a distinction of within- and between-trial variability, which would require either neural recordings (e.g., Steinemann et al., 2022) or specific experimental manipulations (e.g., Ratcliff et al., 2018; Ritz & Shenhav, 2022). Overall, the question to which convenience noise parameters relate to neurocognitive processes remains largely open (Evans et al., 2020; Evans & Wagenmakers, 2019).

Finally, we would like to note that while our analysis indicates that the variance explained by experimental conditions and temporal dependencies are unique, this independence does not provide a rationale for refraining from the inclusion of temporal dependencies in statistical analysis of experimental effects. Incorporating temporal dependencies can still reduce the error variance and can as such be beneficial for statistical detection of the experimental effect (e.g., Baayen & Milin, 2010; Fründ et al., 2014).

### Constraints on generality

The selected datasets in this research reflect the existing literature. As published papers in cognitive neuroscience are highly skewed towards WEIRD (White, Educated, Industrialized, Rich, and Democratic) samples (Henrich et al., 2010), our results are likely skewed in this direction as well. The implications of this bias on the current results remain unclear as cultural differences in cognitive functions are understudied. There is evidence that visual perception can systematically differ between cultures (e.g., in the Müller-Lyer illusion; Heinrich et al., 2010), so we cannot easily state that the more basic functions examined in the current work are safe from selection bias. We are not aware of any empirical evidence on the generality of temporal dependencies, but temporal dependencies seem to have mostly been estimated on WEIRD samples as well. If variability has any biological, social, or evolutionary advantages, it is not implausible that systematic difference in its structures may exist – though this remains an empirical question.

## Conclusion

Experiments are a key tool for identifying factors that influence neural processing and behaviour when we aggregate over trials and/or individuals, but they predict only a fraction of intra-individual variability. Endogenous sources might contribute a much higher proportion to moment-to-moment fluctuations than exogenous influences. In laboratory-derived behaviour data, endogenous or unexplained variability often carries the negative connotation of simply being measurement error. Similarly, in daily life we often focus on reducing variability (e.g., in traffic). However, variability may be an intrinsic and important feature of our biological system (Faisal et al., 2008; Garrett et al., 2013). The larger empirical question is therefore to what extent trial-to-trial variability is predictable. If all possible endogenous sources of variability have been scrutinised, it is possible that some proportion of variability is truly random and thus evades precise trial-to-trial prediction.

## Acknowledgements

We thank the rectorate of Bielefeld University for funding. We are also highly grateful to Aline Bompas, who provided valuable feedback on an earlier version of the manuscript. Finally, we would like to th ank the authors from the original studies for making datasets publicly available.

## Author note

The raw data originated from our own lab is publicly available alongside the analysis code: https://osf.io/pbcus/ Preliminary analyses of this work were presented at Virtual Meeting of the Society for Mathematical Psychology (2021). A previous version of the manuscript has been published on Biorxiv. MNP: conceptualisation, methodology, analysis, investigation, data curation, writing (original draft), and visualisation. TH: conceptualisation, methodology, writing (review & editing), supervision, and funding acquisition. CK: conceptualisation, methodology, resources, validation, writing (review & editing), supervision, and funding acquisition. The authors have no relevant financial or non-financial interests to disclose.

## Supplementary Materials A

Below we provide a brief summary for each of the 30 tasks from which we sourced datasets. More detailed descriptions along with demographics can be found in the original papers.

### Description of congruency-based tasks

1-2. Thirty-eight participants performed a Stroop and a Simon task (Pratte et al., 2010). In the Stroop task, they had to name the colour of the word. The main condition was the written word, which could be congruent, incongruent, or neutral. In the Simon task, participants were shown a red or green square, which they had to press a response button with their right- or left-hand respectively.

The main condition of interest is the side of the presented stimulus (left or right of a central fixation), which could be congruent (e.g., red square presented on the right side of the screen) or incongruent (e.g., red square presented on the left side) to their response hand. For both tasks, participants completed 7 blocks of 74 trials, which were presented with a fixed ITI (700 ms).

3-4. Hundred-and-seven participants took part in two sessions (three weeks apart) in a laboratory experiment, in which they completed the Eriksen Flanker, Stroop, Go/No-Go, and Stop-signal (Hedge et al., 2019). We analysed the data series from the Eriksen Flanker and Stroop tasks, using only the first session.

In the Eriksen Flanker task, participants had to discriminate the direction of a central arrow (left- or rightwards). The main condition for our analysis is the congruency of the four surrounding flankers to the target (congruent, incongruent, or neutral). In the Stroop task, participants had to name the colour of a word. The main condition was the congruency of the written word to the colour (congruent, incongruent, or neutral). For both tasks, participants completed 5 blocks of 144 trials, which were presented with a fixed ITI (1000 ms).

5. Fifty-four participants performed a digit distance task, in which they had respond whether the presented digit was higher or lower than 5 (Rouder et al., 2005). The main condition of interest is the difficulty (distance from 5), which could be easy (2 or 8), medium (3 or 7), or hard (4 or 6). Participants performed 6 blocks of 60 trials, presented at a fixed ITI of 400 ms, during which they received auditory feedback on their accuracy.

6-13. 130 young (18-28 years) and 159 older (64-76 years) participants took part in a two-day experiment involving 11 different neurocognitive tasks. We used the data from two Stroop (Colour and Number) and two Flanker (Arrow and Letter) tasks (Rey-Mermet et al., 2018). Data from the two age groups were analysed separately – thus resulting in 8 separate tasks for our analyses. As per the original paper, participants with high scores on dementia and depression screening questionnaires (Mini-Mental State, Beck Depression Inventory II, Geriatric Depression Scale) were excluded (17 participants), and another two participants were excluded due to experimenter error in the testing session.

In the Colour Stroop task, participants had to name the colour of the word while ignoring the written word, while in the Number Stroop task, they had to respond how many digits were shown, while ignoring the value of the digits. In the Arrow Flanker task, participants had to discriminate the direction of the arrow, while ignoring flankers. In the Letter Flanker task, they had to respond whether the central letter was a vowel (‘S’ or ‘H’) or a consonant (‘E’ or ‘U’), while ignoring the surrounding congruent (e.g., central vowel with vowels), neutral (% or #), or incongruent (e.g., central vowel with consonants) flankers.

For each task, the main condition is congruency (congruent, neutral, or incongruent). Participants completed 4 blocks of 74 trials for each task, presented at a fixed ITI of 1000 ms, during which they were presented with feedback on response accuracy. There was a response deadline of 1900 ms after stimulus onset. All participants were presented with the same stimulus sequence, which contained no complete stimulus repetitions.

14. Thirty-one participants performed a priming congruency task, in which they had to discriminate the left- or rightwards direction of an arrow (Desender et al., 2016). Prior to stimulus onset, a smaller priming arrow was shown for 34 ms, which fitted into the contour of the stimulus arrow (and hence was invisible to the participant), followed by a black screen for 34 ms. If a response was made within 3000 ms, participants were subsequently asked to rate the confidence in their response. The main condition is the congruency of the prime (congruent or incongruent). Participants completed 8 blocks of 80 trials, presented at a fixed ITI of 800 ms.

15. Eighty-six participants performed a priming congruency task (Desender et al., 2014). On each trial, they were first shown a priming arrow for 23 ms (pointed left- or rightwards), which was followed by two masks and a blank screen of 23 ms each. Next, they were presented with a stimulus arrow (160 ms) and were asked to discriminate the direction of the arrow (left or right). The main condition is the congruency of the prime (congruent or incongruent). Participants completed 8 blocks of 60 trials. The ITI was self-paced, and there was a response deadline of 3000 ms after stimulus onset.

16. Thirty participants performed a priming congruency task (Teuchies et al., 2019), in which they had to discriminate the direction of an arrow (left or right), which was shown for 250 ms. Prior to stimulus onset, they were presented with a prime arrow and black screen, both for 17 ms. The prime arrow fitted into the contour of the stimulus arrow and was hence masked. Afterwards, they were asked to rate the confidence in their response.

The main condition is the congruency of the prime (congruent, incongruent, or neutral). Participants completed 6 blocks of 144 trials. The ITI was variable (ranged 1000–5250 ms), and there was a response deadline of 1500 ms after stimulus onset.

### Description of perception-based tasks

1. Fifty-eight participants performed an orientation discrimination task, in which they had to respond whether a grating was oriented left- or rightwards (Rouder et al., 2010). The main condition is the difficulty (tilt of the orientation), which could be easy, medium, or difficult (∼4°, 2°, and 1.5° respectively). Participants completed 9 blocks of 80 trials, which were presented with a fixed ITI (1000 ms), during which they received auditory feedback on their accuracy.

2. Eighteen participants performed a multisensory perception task, in which they had to discriminate the left- or rightwards motion of moving dots (Kayser & Kayser, 2018). An auditory cue was played simultaneously, which could either be congruent or incongruent to the visual information. The coherence of the visual motion (i.e., percentage of dots moving in one direction) could take on four different values (i.e., difficulty levels), which were participant-depended and based on a prior calibration and were adjusted every 35 trials based on the performance. The main conditions are the coherence of the visual motion (total number of percentages is participant-depended) and the audio-visual congruency. All participants performed 5 blocks of 240 trials, except one participant who only performed 4 blocks (960 trials). The stimulus was presented for max 1200 ms (or until response), with a variable trial onset (700-1000 ms) and variable ITI (1500-2000 ms).

3. Twenty-three older participants (aged 62-82) performed the same multisensory perception task as in [2] above (Park, Nannt & Kayser, 2020). Likewise, the main conditions are the coherence of the visual motion (total number of percentages is participant-depended) and the audio-visual congruency. All participants performed 3 blocks of 240 trials.

4-5. Sixteen participants performed two separate auditory discrimination tasks (Kayser et al., 2016). On each trial, they heard two different pure tones presented in background noise. In the frequency task, they were asked which of the tones had the highest frequency. In the intensity task, they were asked which tone was the loudest. Note that the first tone was always fixed (i.e., always the same frequency/loudness), while the second one could vary over seven levels. These levels were participant-dependent and based on a prior calibration session, with the lowest level being 0 (Hz or dB) for everyone. The main condition is the difference in intensity or frequency. Trials were presented at a variable interval (1700-2200 ms) and with a variable duration between the two tones (2400-2562 ms). For the intensity task, all participants performed 4 blocks of 168 (= 672) trials, except one participant who performed 3 blocks (504 trials). For the frequency task, 4 participants performed 3 blocks (504 trials), 1 participant performed 5 blocks (840 trials), and 11 participants performed 4 blocks (672 trials).

6. Thirty participants performed a detection task, featuring auditory, visual, and audio-visual targets (unpublished pilot data). The target was presented at the left or the right side, and participants had to detect it. The main condition is stimulus type. All participants completed 4 blocks, with block size being 239 trials for 7 participants (= 956 trials), and 215 trials for the others (860 trials).

7. Thirty participants performed a noisy detection task (Maniscalco et al., 2017). On each trial, they were shown two patches of noise for 33 ms, and were asked which patch contained a target. The target’s contrast was fixed and participant-depended (based on a 120-trial calibration). Afterwards, participants rated the confidence in their response. The main condition of interest is the location of the target (left or right). Participants performed 10 blocks of 100 trials, presented at a fixed ITI of 1000 ms.

8. Forty-one participants performed a noisy detection task, with an interleaved memory task (Maniscalco et al., 2017). We analyse the data from the detection task only. On each trial, they were shown two patches of noise for 33 ms, and were asked which patch contained a target. The target’s contrast was presented at three different difficulties (participant-depended, based on calibration). Afterwards, participants rated the confidence in their response. The main conditions are the location (left or right) and the contrast of the target. Participants performed 5 blocks of 102 trials, collected over two days. Trials were presented with a fixed ITI of 1000 ms.

9. Fifteen participants performed an orientation discrimination task (Samaha et al., 2016), in which they had respond whether a grating (shown for 33 ms) was titled left- or rightwards at 45°. The main condition is the difficulty (easy or hard), which related to the noise of the grating, and was participant-depended (based on prior calibration). Participants performed 2 blocks of 160 trials, collected over two days. Trials were presented at a variable ITI (300-500 ms).

10-12. Twenty participants performed three tasks each (Samaha et al., 2017). In the contrast perception task, participants were shown two gratings on each trial, and were asked which grating had the highest contrast. In the orientation discrimination task, participants were also shown two gratings, and had to respond whether their orientations were same or different (25° more tilt). In the visual short-range memory task, participants were shown one target, followed by a 3 s blank screen, and then the second target. Likewise, they had to respond if the orientations were the same or different.

For all tasks, the main condition is the contrast of the stimuli. Participants performed 3 blocks of 100 trials for each task, which were presented at a variable ITI (300-500 ms). Only for the contrast perception task, one participant only completed 2 blocks, and one participant only 1 block – the latter was excluded from analysis on this task. After each trial, participants had to rate the confidence in their response. The contrast was individually determined with a prior calibration and adaptive throughout each task.

13. Twenty-four participants performed a colour judgment task (Desender et al., 2019). On each trial, they were shown 8 coloured dots presented in a circle around a central fixation. Participants had to judge whether the average colour across all dots was bluer or more red. Next, they were asked to rate the confidence in their response (though half of the participants only were asked for a rating in the even blocks). The main conditions of interest are the mean and the variance of the colour. Half of the participants performed 8 blocks of 60 (= 420) trials, and half performed 8 blocks of 64 (= 512) trials. Trials were presented with a fixed ITI of 1000 ms, and there was a response deadline of 1500 ms.

14. Twenty-one participants performed an emotion discrimination task, in which they had to respond whether the presented face was angry or happy (Johannknecht & Kayser, 2022). The stimulus face was shown for 100 ms. The main condition is the emotion (angry/happy). Participants performed 2 blocks of 200 trials, with trials being presented with a variable trial onset 400-1000 ms) and ITI (1200-1500 ms).

### Model fitting

Each dataset was fitted with a GLM. The model for datasets with one experimental manipulation was fitted as:

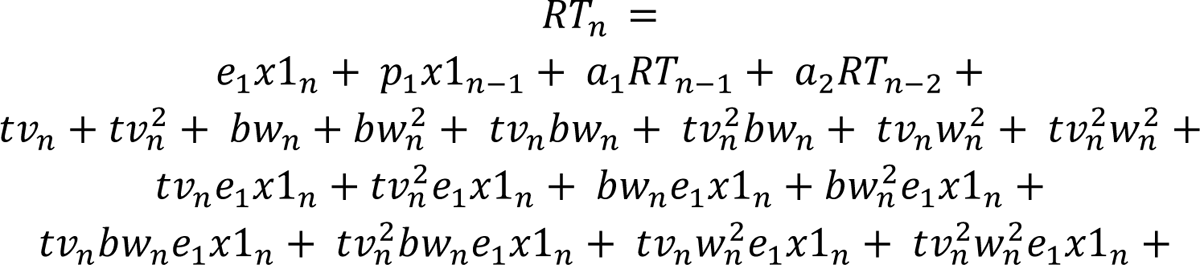

and the fitted model for datasets with two experimental conditions is:

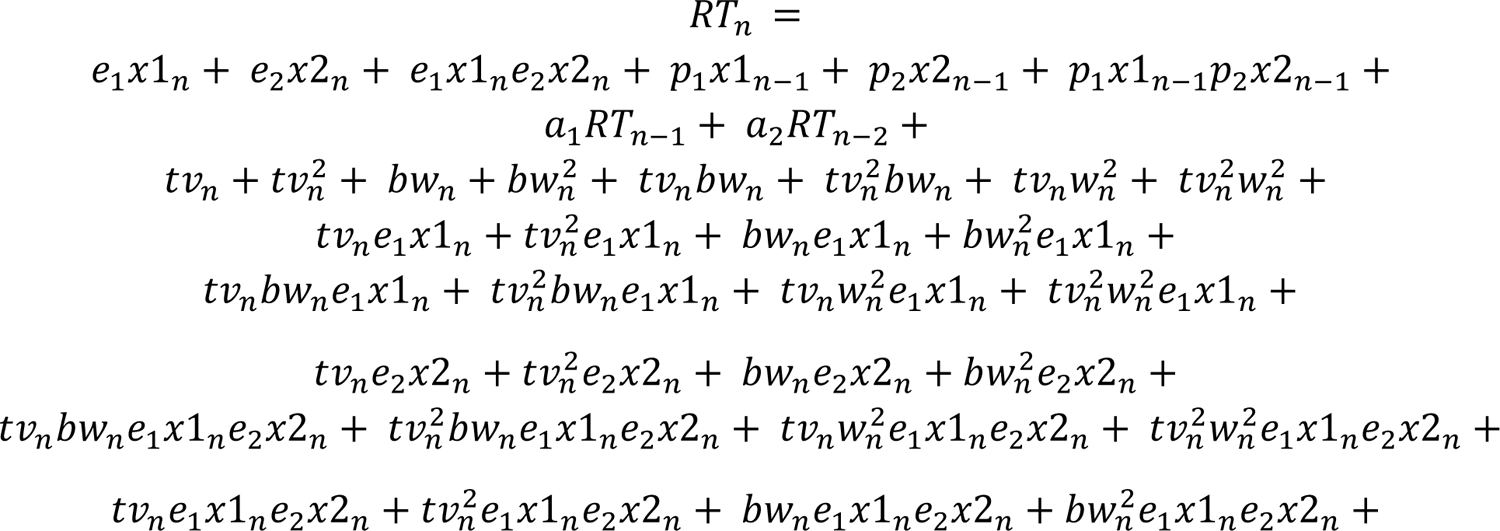

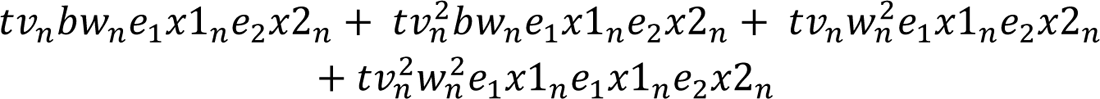

with e_1_ and e_2_ reflecting the weights for both of the experimental manipulations *e* multiplied respectively by observation *x1* and *x2* on trial *n,* p_1_ and p_2_ reflecting the weights of the experimental manipulations from the previous trial *n-1*, a_1_ and a_2_ reflecting the weights for the relation with the RT from trial *n-1* and *n-2*, and *t* reflecting the weight of trial number multiplied by the current blockwise trial number *v*, and *b* reflecting the weight of block number multiplied by the current block number *w.* These models include main effects for all predictors, as well as each possible interaction between current condition, trial number, and block number.

## Supplementary Materials B

**Figure B1.**
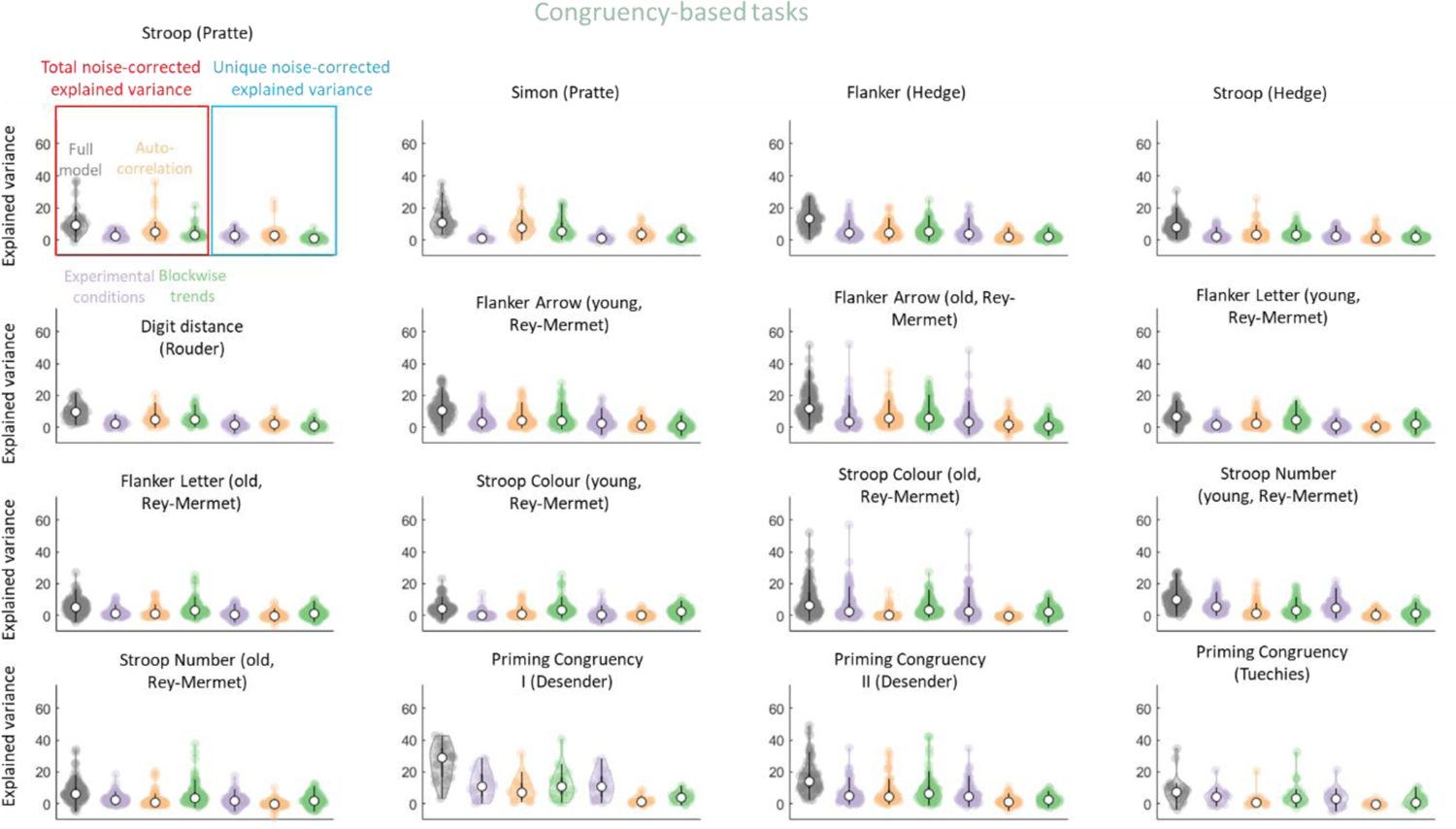
Percentage of noise-corrected total explained variance and unique explained variance in congruency-based tasks. Each panel shows one task, with the task name and first author name on top. In each panel, each dot represents one dataset, the white circle represents the median of the distribution, and the width of the dot cloud indicates distribution density – by the three sources combined (full model or R^2^_abc_; grey) as well as separately (purple, orange, and green for experimental condition R^2^_c_, autocorrelational R^2^_a_, and blockwise trends R^2^_b_ sources respectively) as estimated by the dataset level GLMs. The first four violins show the total explained variance (red square) and the last three show the unique explained variance (light blue square). Negative values indicate instances in which the predictors explained less variance than random noise.

**Figure B2.**
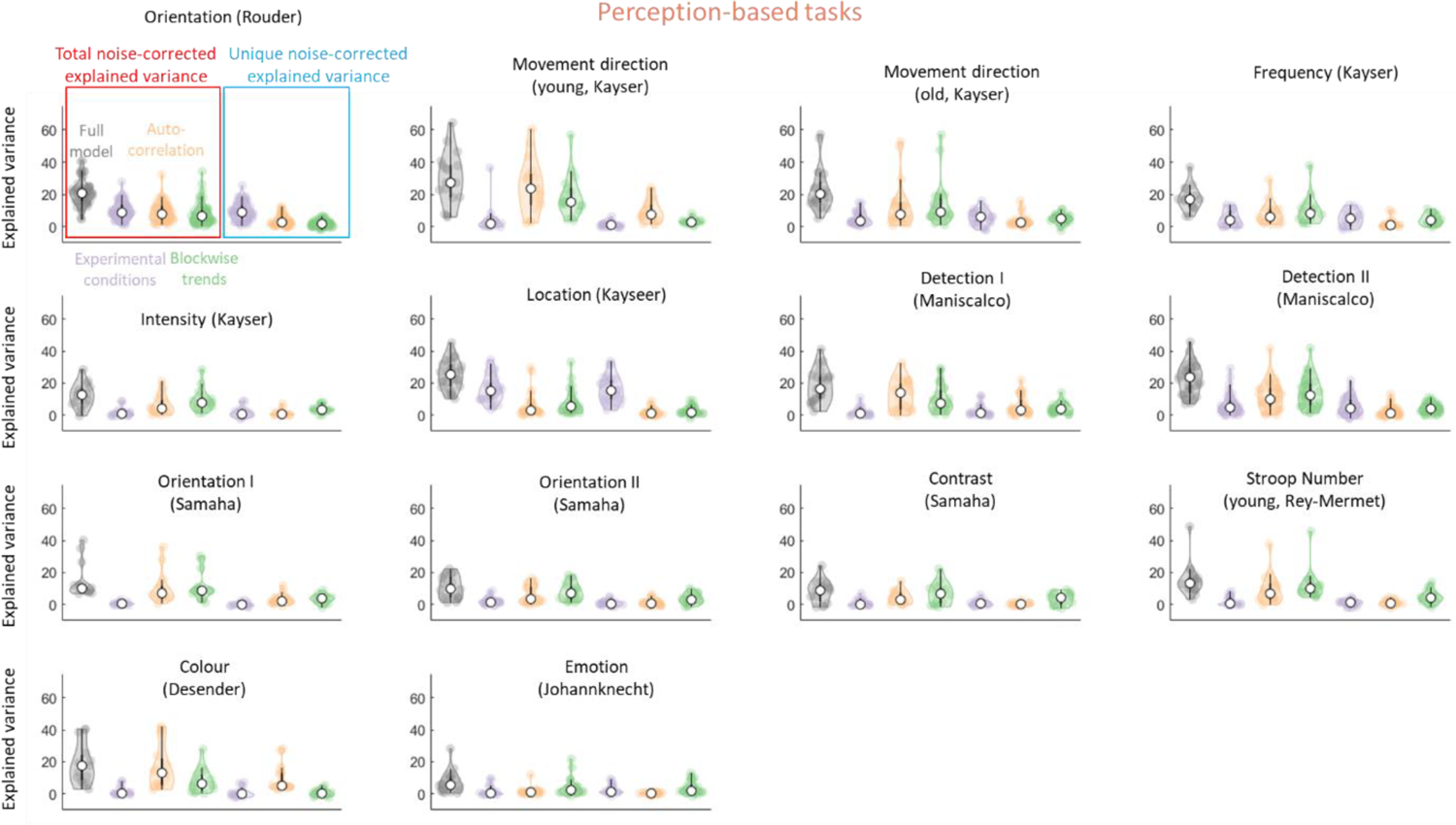
Percentage of noise-corrected total explained variance and unique explained variance in perception-based tasks. Conventions are the same as in Figure B1.

## Supplementary Materials C

Here we report the results of various control analyses that were conducted to check the robustness of our result patterns. We check for the effect of different analysis choices, additional systematic sources of variance not implemented in the current models, impact of multicollinearity, and impact of experimental condition and blockwise trends on the autocorrelation. Congruency- and perception-based tasks were combined for these control analyses, based on the consistent patterns we had observed across both task types.

Because of the number of observations per distribution, inferential tests like paired t-tests can likely detect very small but meaningless differences – and thus do not inform about the robustness of the result patterns. Therefore, we compare the median values of explained variance between our original and control analysis. Any differences < 1% were considered as not noteworthy.

**Table C1.**
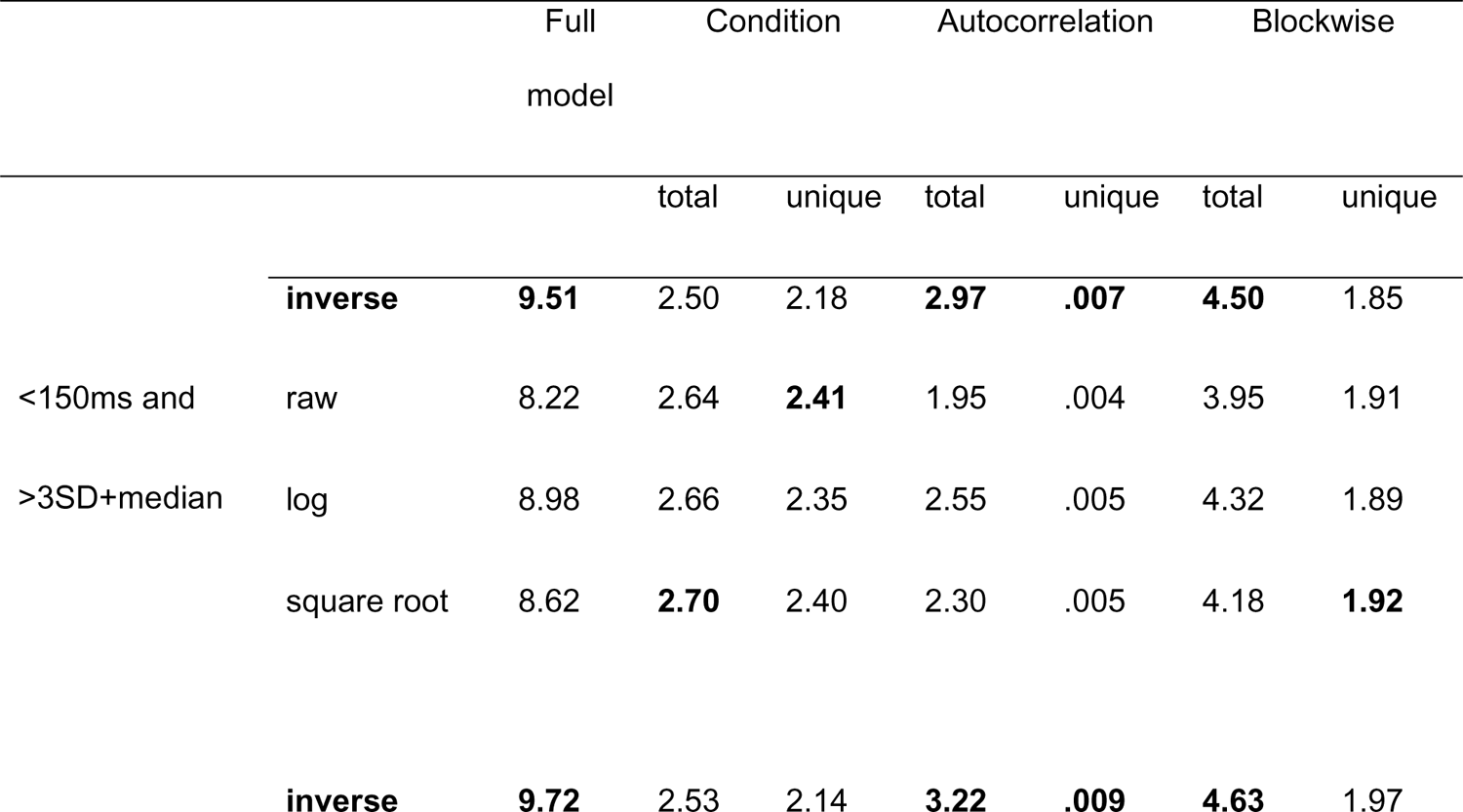
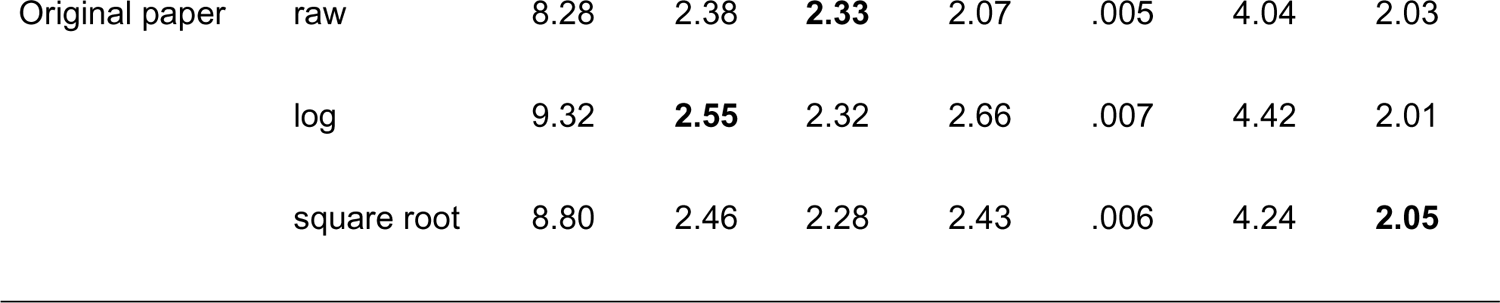
Median percentage of noise-corrected explained variance across all participants and all tasks for the full model (R^2^_abc_) and the total plus unique explained variance for each of three sources separately under (2 outlier exclusion x 4 data transformations =) 8 different analysis choices. The numerically highest value for each R^2^ is shown in bold.

|  |  | Full<br>model | Condition |  | Autocorrelation |  | Blockwise |  |
| --- | --- | --- | --- | --- | --- | --- | --- | --- |
|  |  |  | total | unique | total | unique | total | unique |
|  | **inverse** | **9.51** | 2.50 | 2.18 | **2.97** | **.007** | **4.50** | 1.85 |
| <150ms and | raw | 8.22 | 2.64 | **2.41** | 1.95 | .004 | 3.95 | 1.91 |
| >3SD+median | log | 8.98 | 2.66 | 2.35 | 2.55 | .005 | 4.32 | 1.89 |
|  | square root | 8.62 | **2.70** | 2.40 | 2.30 | .005 | 4.18 | **1.92** |
|  | **inverse** | **9.72** | 2.53 | 2.14 | **3.22** | **.009** | **4.63** | 1.97 |
| Original paper | raw | 8.28 | 2.38 | **2.33** | 2.07 | .005 | 4.04 | 2.03 |
|  | log | 9.32 | **2.55** | 2.32 | 2.66 | .007 | 4.42 | 2.01 |
|  | square root | 8.80 | 2.46 | 2.28 | 2.43 | .006 | 4.24 | **2.05** |

### Effect of analysis choices

Here, we ask how our results are affected by preprocessing choices. In our main analyses, the RT series in each dataset were cleaned and transformed prior to analysis. RT outliers were removed with one common criterion for outlier exclusion across all participants: a lower cut-off of 150 ms and an upper cut-off of 3 SD above the median RT for outlier exclusion. Next, RT series were inverse-transformed, and then the GLMs were run on to predict to RT series and obtain the measures of explained variance (as shown in Figure 5). The first results row of Table C1 shows the explained variance by our original analyses by taking the median across all datasets.

Here we instead use logarithmic (natural log) and square root to transform the RT series prior to analysis. Likewise, GLMs were run on to obtain the measures of explained variance, and the median explained variance was calculated for each source. GLMs were also run on the raw RT series (i.e., no transformation at all). This allows for comparison of the median explained variance across four different transformation choices.

We also checked for the effect of outlier exclusion. The RT exclusion method was taken from the original paper if available (reported in Table 2 and 3). Note that some papers only analysed accuracy and therefore did not include an RT exclusion method. For these papers we kept the < 150 ms and > median RT plus 3 SD(RT) cut-off. Again, analyses were run using all four transformation methods (Table C1; bottom).

For each transformation separately, the total and unique explained variance was extracted for each dataset. We obtained qualitatively very similar results across all analysis choices (see Table B1 for the median explained variance across all participants and tasks). GLMs with raw RT series explained the least amount of variance, but the differences between inverse, log, and square root transformations in total explained variance were all lower than 1%.

### Additional sources of variance

Our analyses were focused on the experimental conditions of interest. However, the analysed tasks contain additional exogenous sources, such as a variable ITI or a redundant visual feature of the stimulus (e.g., the actual direction of the target arrow in the Flanker task). On the group level, the effects of arrow direction are often not statistically different from zero, but for each individual participant, the explained variance will numerically be higher than zero. As a control analysis, we therefore ran single-subject GLMs to examine if the amount of explained variance substantially increases when taking additional exogenous sources into account.

GLMs were computed for all datasets that included additional exogenous sources that were uncorrelated with the experimental conditions, if made available at their archival source (11 datasets in total across congruency and perception-based tasks). For example, in the Simon task from Pratte et al. (2010), the colour and location of the stimulus varied across the trials independently from the experimental condition. Both of these factors were added to the new GLM. In this particular example, the additional factors explained .007% of the total variance. It should be noted that the interpretation of these sources can be difficult. For example, in the flanker task, participant respond to right arrows with their right hand and to left arrows with their left hand. This means that any additional explained variance could be due to the visual stimulus property, the motor response bias (i.e., being faster with dominant hand), or a combination of both. We therefore do not make strong claims about what causes this increase in explained variance, but only note that it is relatively small.

Table C2 shows an overview of the additional exogenous factors added to the GLM, as well as the original explained variance by the full model (Figure 3 and 5) and the extended model’s explained variance. Overall, increases in explained variance were small, with a median increase of 1.16%.

**Table C2.**
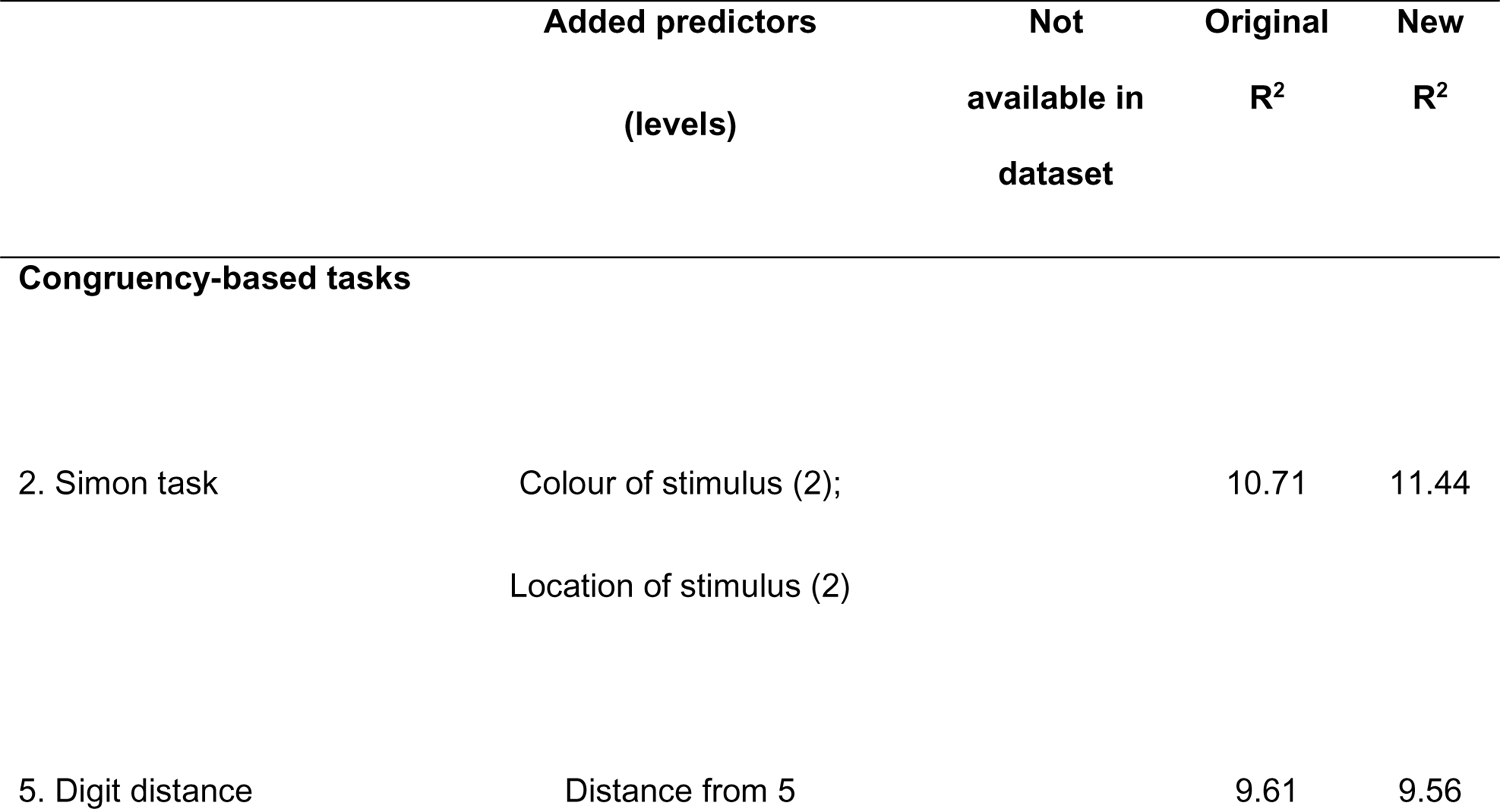
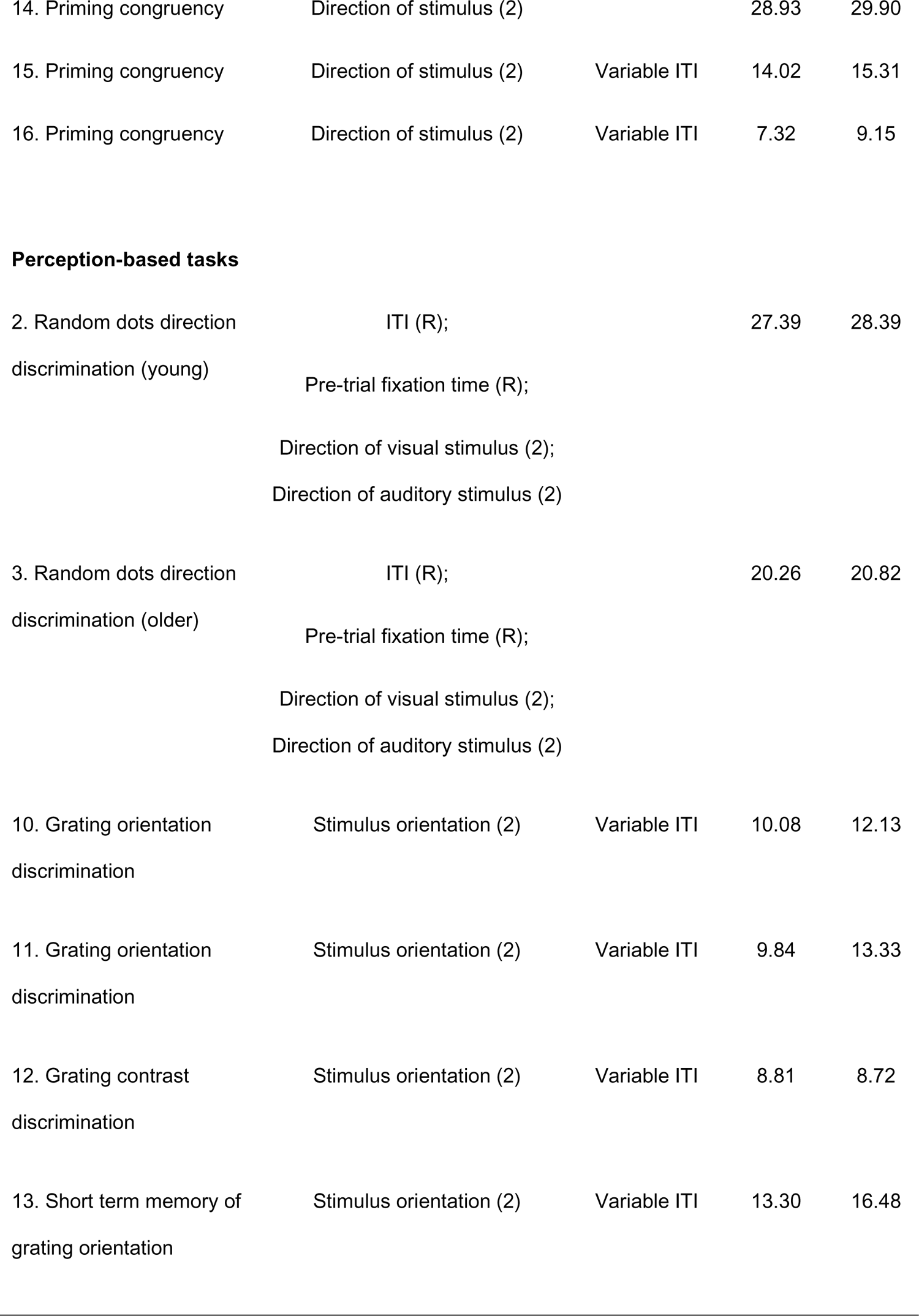
Overview of analysed datasets in GLMs featuring additional exogenous sources. Shown are the 11 included task, with the exogenous sources that have been added to the new GLM (and the exogenous sources that could not be included due to non-availability in the datasets, where applicable), with their original and new amount of noise-corrected explained variance (R^2^). Increases in explained variance are minimum for most of the datasets.

| | Added predictors<br>(levels) | Not<br>available in<br>dataset | Original<br>$R^2$ | New<br>$R^2$ |
| --- | --- | --- | --- | --- |
| **Congruency-based tasks** |  |  |  |  |
| 2. Simon task | Colour of stimulus (2);<br>Location of stimulus (2) |  | 10.71 | 11.44 |
| 5. Digit distance | Distance from 5 |  | 9.61 | 9.56 |
| 14. Priming congruency | Direction of stimulus (2) |  | 28.93 | 29.90 |
| 15. Priming congruency | Direction of stimulus (2) | Variable ITI | 14.02 | 15.31 |
| 16. Priming congruency | Direction of stimulus (2) | Variable ITI | 7.32 | 9.15 |
| Perception-based tasks |  |  |  |  |
| 2. Random dots direction<br>discrimination (young) | ITI (R);<br>Pre-trial fixation time (R);<br>Direction of visual stimulus (2);<br>Direction of auditory stimulus (2) |  | 27.39 | 28.39 |
| 3. Random dots direction<br>discrimination (older) | ITI (R);<br>Pre-trial fixation time (R);<br>Direction of visual stimulus (2);<br>Direction of auditory stimulus (2) |  | 20.26 | 20.82 |
| 10. Grating orientation<br>discrimination | Stimulus orientation (2) | Variable ITI | 10.08 | 12.13 |
| 11. Grating orientation<br>discrimination | Stimulus orientation (2) | Variable ITI | 9.84 | 13.33 |
| 12. Grating contrast<br>discrimination | Stimulus orientation (2) | Variable ITI | 8.81 | 8.72 |
| 13. Short term memory of<br>grating orientation | Stimulus orientation (2) | Variable ITI | 13.30 | 16.48 |

### Multicollinearity

As our models feature a large number of predictors, we tested if these predictors met the assumption of absence of multicollinearity. For each participant, the tolerance value and Variance Inflation Factor were calculated for each predictor. There is reason to assume severe multicollinearity for 8 of the 31 datasets, defined as a tolerance > .10 and a VIF > 10, for the experimental predictors for almost all participants. These were tasks that either featured two experimental conditions and/or a stimulus that changes throughout the task (e.g., stimulus contrast adjusting to current performance), and were all perception-based tasks. We removed these 8 tasks from the dataset and assessed whether the overall result pattern would be affected; such a change of results would imply that our analysis and conclusions are contaminated by predictor multicollinearity. However, the amount of explained variance for the different variability sources was similar with and without the 8 identified tasks: the total explained variance dropped by less than 1%, and by .001-.39% for each separate source (see Table C3), suggesting that multicollinearity did not impact our conclusions.

**Table C3.**
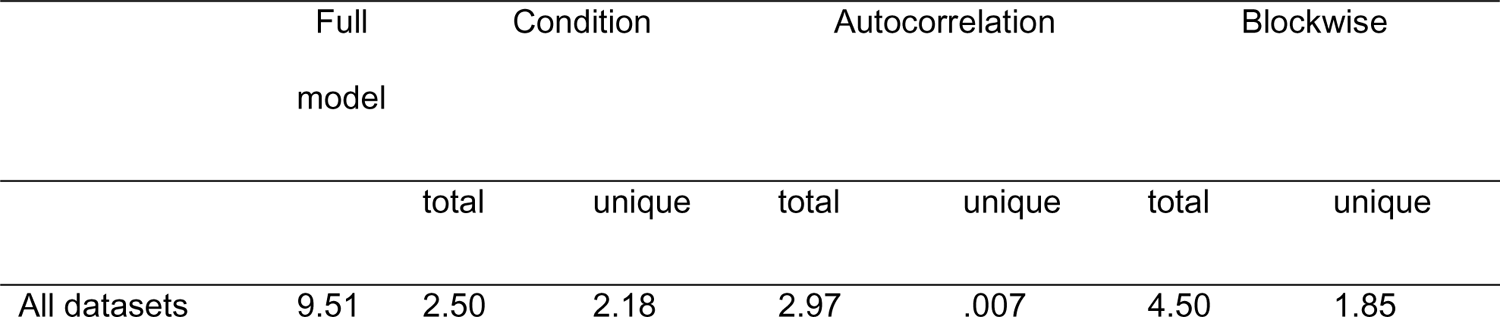
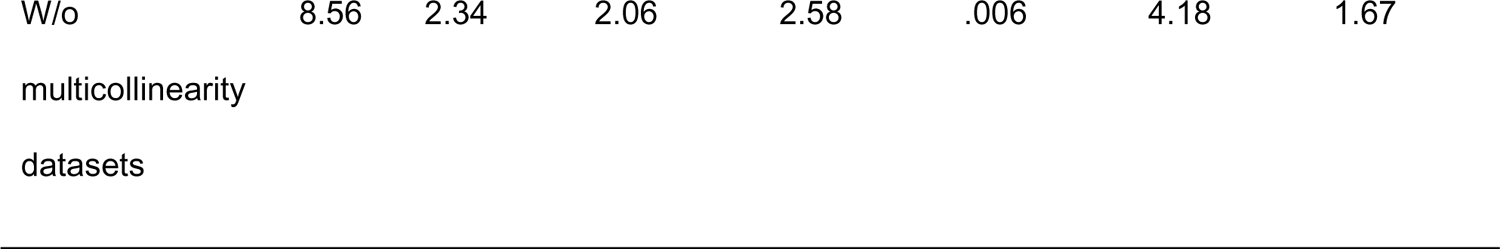
Median percentage of noise-corrected explained variance across all participants for the full model (R^2^_abc_) and the total plus unique explained variance, after excluding the datasets with indications of serious multicollinearity.

### Correcting the autocorrelation for the other predictors

In our main analyses, we first estimate our three sources separately and combined, and then calculate the communalities and unique explained variance. Alternatively, one may employ a sequential approach. Here, RT was first regressed on the experimental conditions and blockwise trends. We then obtain the previous RT on trial *n* and *n-1* from the residuals, and run the full model with these residuals, experimental conditions and blockwise trends. Again, the amount of explained variance for was similar, differing only .2% in explained variance (see Table C4).

**Table C4.**
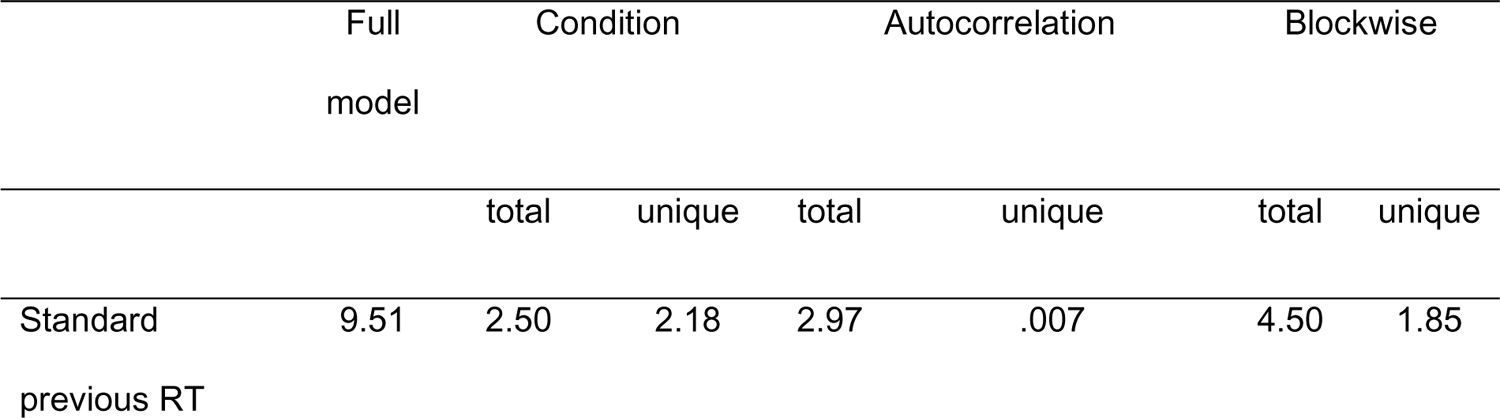
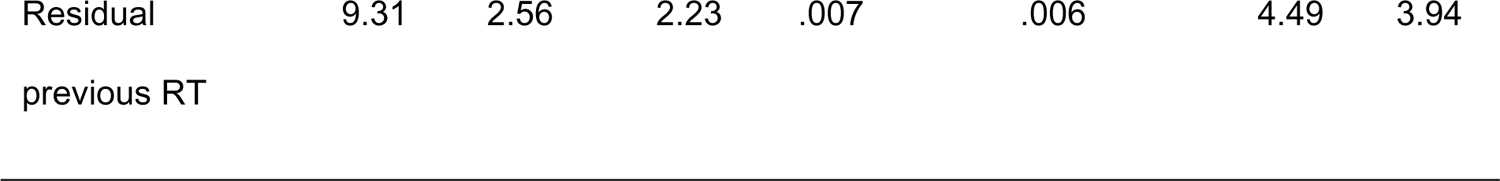
Median percentage of noise-corrected explained variance across all participants for the full model (R^2^_abc_) and the total plus unique explained variance, when first regressing the experimental conditions and blockwise trends from the previous RT series before calculating the models.

## Supplementary Materials D

Linear Mixed Models (LMM) explain variance as parts common to all instances of a grouping variable (i.e., fixed effects) and the variability of these common parts across the grouping variable (i.e., random effects). The more random effects are included in the model, the more data and computation time are required for model fitting. Moreover, correlations between predictors further increase the required amount of data for fitting. For our purposes, we ideally would want to assign all model terms to both the fixed and the random effects, effectively allowing the model to identify variability of all factors and interactions of our model that is common to all participants, but also to determine the variability between them for all fixed model terms. This model did not run successfully due to being too unstable. We therefore chose the common strategy of entering all model terms in the fixed effects while making the random effects increasingly complex.

With this approach, the fixed factor structure is identical to that of our GLMs. The difference between our GLM approach and an LMM approach is that we fit a separate GLM for each dataset and inspect the resulting amounts of explained variance, whereas the LMM fits all participants at once and assigns variability to the fixed and random model parameters. Thus, the information provided between the GLM and LMM approaches is different, because there is no equivalent of the random factor structure in our GLM approach. Thus, the LMM assigns some part of the variance to differences between participants and might, thus, explain more variance overall than the GLM. However, this additional explained variance does not address the question we posed in our paper, namely how well single trial RT data can be predicted; instead, it quantifies variability between participants.

We explored two different LMM approaches. First, we conducted LMMs on the RT series across all datasets separately for congruency- and perception-based tasks. Next, we conducted LMMs on the RT series across all datasets separately for each task.

### All datasets

LMMs were run in R (R Core Team, 2013) with the lmer function in the lme4 package (Bates et al., 2015) for each of the three sources separately as well as combined (full model), mimicking the GLM analyses presented throughout the paper. For each source, three models were run with increasingly complex random terms, including: 1) only the intercept, 2) the intercept plus slope A, and 3) the intercept plus slope A plus slope B. As the more complex models took several hours to days to runs and did not converge reliably, we did not add any more terms. The random effects were allowed to vary over unique participant number (note, rather than by dataset) and task. This strategy takes into account when a participant contributed data to several experiments for estimating model parameters. First, for experimental condition, slope A was the experimental condition of the current trial, and slope B was the condition of the previous trial – giving us the following three random effects:

– *(1 | participant) + (1 | task)*

– *(1 + condition | participant) + (1 + condition | task)*

– *(1 + condition + previous condition | participant) + (1 + condition + previous condition | task)*

In Figure D1, these are indicated by the red triangle, blue plus, and green star respectively. Secondly, for the autocorrelation, we included RT-1 as slope A and RT-2 as slope B. Thirdly, for the blockwise trends model, we included trial number as slope A and block number as slope B. The most complex models (with slope A and B) frequently did not converge and once did not successfully run at all (blockwise trends model) – indicating that even the large amounts of data are insufficient to fit them. For the full model, we included condition and previous condition as slope A and B, because these are the simplest terms. Because the most complex model again had severe convergence problems, we did not include any of the other model terms.

**Figure D1.**
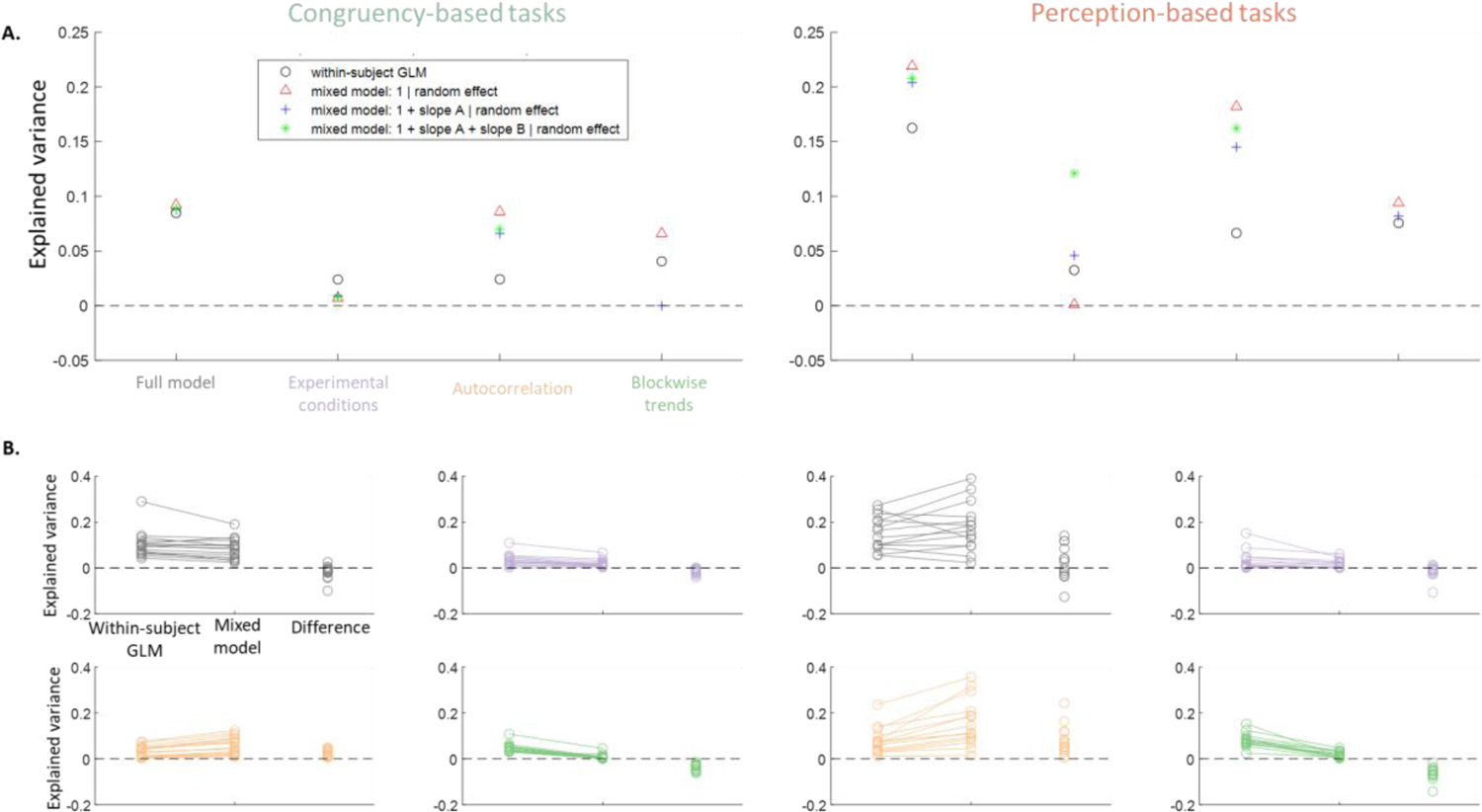
Comparison of the within-subject GLM analyses and across-subject LMM analyses. The percentage of noise-corrected total explained variance is shown for the full model (grey), experimental conditions model (purple), autocorrelation model (orange), and blockwise trends (green) model. **A.** Median explained variance by the GLM (black circle) and explained variance by three LMMs with increasing complexity (red triangle, blue plus, and green star respectively), separately for congruency- (left) and perception-based tasks. **B.** Median explained variance by the GLM and explained variance by the LMM, with each line representing one task. The right circles indicate the difference between the GLM and LMM, with difference values above zero indicating that the GLM explained more variance.

Estimates of explained variance was extracted for each model using the MuMIn package (Johnson, 2014; Nakawaga et al., 2017; Nakagawa & Schielzeth, 2013). For the congruency-based tasks, datasets from all 16 tasks were run in one model. For the perception-based tasks, this was not possible, because 7 included one experimental condition while the other 7 included two (see Table 2). We therefore ran LMMs separately for one- and two-condition tasks, and then calculated the mean across both. As with the GLM, we performed a noise correction procedure by estimating the explained variance on 100 shuffled RT series and subtracting the mean from the original estimates – though in contrast to the GLM, these values were near zero, as the LMM is more protective against overfitting. These corrected estimates were plotted in Figure D1A, alongside the median explained variance across datasets by the GLMs (as reported in Table 3). The total explained variance across the four models was similar, with all symbols nearly overlapping. In contrast, the experimental conditions explained more variance in the GLM, while both temporal dependencies explained more variance in the LMM.

### Datasets per task

Next, we ran LMMs separately for each task. Again, we chose to run each model with increasing complexity, including intercept, slope A, and slope B. Here, the random model terms can only vary over participants, rather than over participants and tasks. For example, for the experimental condition model, we fit the random terms:

– *(1 | participant)*

– *(1 + condition | participant)*

– *(1 + condition + previous condition | participant)*

Again, the more complex models did not converge reliably or did not fit successfully. Furthermore, adding more random model terms did not seem to impact the explained variance, with differences between the levels of complexity often < .001. Here, we therefore only include the models with a random intercept, which converged well. Again, all estimates of explained variance were corrected for noise. Figure D1B shows median explained variance per task by the GLM and the explained variance by the LMM alongside their difference, for the full model and each source separately.

